# High levels of the cardiomyocyte-specific kinase Tnni3k impair zebrafish heart regeneration by driving chronic myocardial inflammation

**DOI:** 10.64898/2026.09.03.749026

**Authors:** Miriam Fernández-Lajarín, Sean Keeley, Juan Manuel González-Rosa

## Abstract

**Background:** Zebrafish regenerate their hearts after injury, and defining the barriers that block this capacity in mammals may reveal targets for heart failure treatment. Elevated levels of the cardiomyocyte-specific kinase TNNI3K are associated with human cardiomyopathy, and its overexpression drives adverse remodeling in mice. Recent work has linked elevated TNNI3K to cardiomyocyte polyploidization and loss of regenerative competence, but whether this cell-cycle effect accounts for the pathology seen in patients has not been established. Because TNNI3K is restricted to cardiomyocytes, whether it also acts non-cell-autonomously is unknown.

**Methods:** We generated an allelic series of zebrafish lines comprising cardiomyocyte-specific Tnni3k overexpression, an expression-matched kinase-dead variant, a full-locus deletion, and a line overexpressing mouse Tnni3k. Regeneration, cardiomyocyte proliferation, and ploidy were assessed in uninjured hearts and after cryoinjury. We combined RNA-sequencing, macrophage depletion before or after injury, tamoxifen-inducible Cre-lox switches, and cardiomyocyte-restricted CRISPR mutagenesis. Hearts from mice overexpressing human TNNI3K were analyzed in parallel.

**Results:** Elevated Tnni3k impaired heart regeneration in a kinase-dependent manner. Cardiomyocyte proliferation, redifferentiation, and productive cell division were preserved, and polyploidization remained well below the threshold compatible with robust regeneration, arguing against a cell-cycle defect. Instead, Tnni3k overexpression established an inflammatory and fibrotic state in the uninjured myocardium, marked by interferon, NF-κB, and antigen-presentation programs and by leukocyte and macrophage accumulation. This state amplified after injury, and these animals retained substantially more scar at 60 days post-injury. Depleting macrophages before injury, but not after, prevented the excess fibrosis. Switching Tnni3k off after injury attenuated both inflammation and fibrosis. Cardiomyocyte-specific deletion of *pkmb*, a glycolytic gene strongly repressed in Tnni3k-overexpressing animals and previously associated with inflammation, reproduced both phenotypes. Mice overexpressing TNNI3K showed comparable fibrosis and macrophage accumulation.

**Conclusions:** Elevated Tnni3k impairs cardiac regeneration by sustaining a chronic inflammatory and fibrotic state rather than by driving polyploidization. Because this state requires continued kinase activity and remains reversible after injury, TNNI3K inhibition may warrant exploration as a strategy to interrupt the inflammation-fibrosis loop in inflammation-driven cardiac disease.

**What is New?:**

- Elevated Tnni3k (troponin I-interacting kinase) impairs heart regeneration in zebrafish without reducing cardiomyocyte proliferation or productive cell division, and with only marginal polyploidization, arguing against the prevailing ploidy-based model.
- Cardiomyocyte-specific Tnni3k overexpression establishes an inflammatory and fibrotic state in the uninjured heart that is amplified after injury and fails to resolve; depleting macrophages before injury, but not after, prevents the excess fibrosis.
- Switching Tnni3k overexpression off after injury attenuates inflammation and fibrosis. Cardiomyocyte-specific mutagenesis identifies the downregulation of pyruvate kinase (*pkmb*) as a candidate downstream effector linking Tnni3k to innate immune activation.

**What are the Clinical Implications?:**

- Chronic myocardial inflammation can act as a barrier to cardiac repair independently of cardiomyocyte cell-cycle activity, suggesting that strategies aimed solely at stimulating cardiomyocyte proliferation may be insufficient in hearts carrying a pre-existing inflammatory substrate.
- Because elevated Tnni3k primes the myocardium before injury, patients with increased TNNI3K expression or gain-of-function variants might enter an ischemic event with an inflammatory substrate already in place.
- Since the phenotype remains reversible after injury and selective TNNI3K inhibitors already exist, post-injury TNNI3K inhibition could potentially be explored to limit fibrosis in cardiac disease driven by persistent inflammation.

## INTRODUCTION

Adult mammals fail to regenerate their hearts after injury. Following myocardial infarction, lost muscle is replaced by a fibrotic scar rather than by cardiomyocytes. In contrast, zebrafish regenerate their hearts robustly throughout life^1–3^. Work over the last two decades has identified specific differences between zebrafish and adult mammals that explain the differences in their regenerative abilities. Zebrafish and other animals capable of regenerating their hearts have predominantly diploid cardiomyocytes that re-enter the cell cycle and successfully divide, completing cytokinesis^2,4^. However, cellular proliferation alone is insufficient; successful regeneration also requires a permissive microenvironment, including the formation of a transient scar and a tightly regulated, self-resolving inflammatory response. In mammals, a prolonged and unresolved inflammatory response after infarction is a recognized driver of adverse remodeling, excessive scarring, and heart failure^5,6^.

Although it has been widely accepted that cardiac injury inevitably results in permanent fibrosis and a decline in heart function in mammals, recent findings suggest that the heart’s response to injury is a variable trait influenced by several genes^7,8^. One potential regulator of this outcome is the cardiac-specific troponin I-interacting kinase (*Tnni3k*). The association between elevated Tnni3k and cardiac pathology is well established in both animal models and humans. *TNNI3K* is among the most strongly upregulated genes in samples from human hearts with end-stage dilated cardiomyopathy^9,10^ and ischemic cardiomyopathy^11^. Moreover, *TNNI3K* variants have been associated with a wide spectrum of cardiac diseases, including conduction disease, dilated cardiomyopathy, supraventricular arrhythmia, and sudden death^12–15^. Large-cohort burden testing indicates that the dilated-cardiomyopathy signal is driven by missense variants that increase kinase activity, whereas loss-of-function alleles are tolerated^16^. Consistent with human data, elevating Tnni3k in mouse models accelerates disease progression, adverse remodeling, and fibrosis under various stresses^17^, whereas its genetic ablation is protective^18^.

Despite its clear clinical relevance, the mechanisms by which this cardiac-specific kinase drives disease remain debated, with recent research focusing primarily on its role in cell-cycle regulation. Initial studies in mice linked elevated Tnni3k levels to increased cardiomyocyte polyploidization and loss of proliferative capacity, leading to the hypothesis that Tnni3k overexpression impairs regeneration by driving myocardial polyploidization^8^. However, this interpretation has recently been challenged. Reuter and colleagues demonstrated that elevated Tnni3k promotes cardiomyocyte S-phase entry after injury, but that these events ultimately lead to polyploidization rather than cell division^19^. Complementing these findings, Purdy and colleagues showed that loss of Tnni3k reduces S-phase entry yet paradoxically increases the number of cardiomyocytes that successfully complete cell division^20^. Together, these studies support a model in which Tnni3k drives S-phase entry while suppressing cytokinesis in the mammalian heart. However, it remains unknown whether these alterations in ploidy mechanistically explain the adverse remodeling and clinical pathologies associated with elevated Tnni3k, or if other disease-driving mechanisms are at play.

This leaves two critical gaps in our understanding. First, to disentangle the kinase’s role in proliferation from the intrinsic mammalian block to cell division, it is necessary to study how Tnni3k functions in a naturally regenerative species in which cardiomyocytes are fully competent to divide. Second, because Tnni3k is selectively expressed in cardiomyocytes, it remains completely unexplored how its elevation drives the pathology seen in patients and animal models, and whether it triggers non-cell-autonomous programs independent of its cell-cycle effects.

Here, we define the interplay between Tnni3k, myocardial regeneration, and inflammation in adult zebrafish. We show that *tnni3k* is broadly expressed in the myocardium and is upregulated after cryoinjury. Using a new allelic series of gain- and loss-of-function models, we find that elevated Tnni3k blocks regeneration in a kinase-dependent manner. Strikingly, this regenerative failure is not driven by cardiomyocyte polyploidization, which remains well below pathogenic thresholds. Instead, sustained Tnni3k overexpression establishes a pre-injury inflammatory state that amplifies after injury, leading to excessive, unresolved fibrosis. Using macrophage ablation and temporal genetic manipulations, we identify this chronic inflammatory environment and the continuous requirement for Tnni3k kinase activity as the primary drivers of failed regeneration. Finally, we show that myocardial deletion of the pyruvate kinase *pkmb*, one of the genes most strongly repressed in response to Tnni3k overexpression, mimics the inflammatory and fibrotic phenotypes. Together, our results redefine Tnni3k as a central regulator of the cardiac microenvironment and a potential therapeutic target for inflammation-driven heart failure.

## RESULTS

### *tnni3k* expression is upregulated following injury, and its overexpression drives ventricular enlargement with only modest effects on cardiomyocyte ploidy

Tnni3k has been associated with cardiomyocyte polyploidy, but whether it is naturally expressed in species with hearts composed predominantly of diploid cardiomyocytes remains unknown. The zebrafish genome contains a highly conserved copy of *tnni3k* on chromosome 8, whereas only ∼1-2% of all cardiomyocytes are naturally polyploid^4^. Zebrafish Tnni3k contains the same domains described in human TNNI3K, and all residues previously identified as modifiers of the kinase activity^21^ are conserved **(Fig. S1A-S1B)**. To test whether *tnni3k* expression is restricted to the rare polyploid fraction, we performed RNAscope *in situ* hybridization on sections of adult hearts and detected widespread myocardial expression of *tnni3k* in both the atrium and the ventricle **(Fig. 1A,B)**. Thus, contrary to our initial hypothesis, *tnni3k* is broadly expressed by cardiomyocytes rather than being limited to the polyploid fraction.

**Figure 1.**
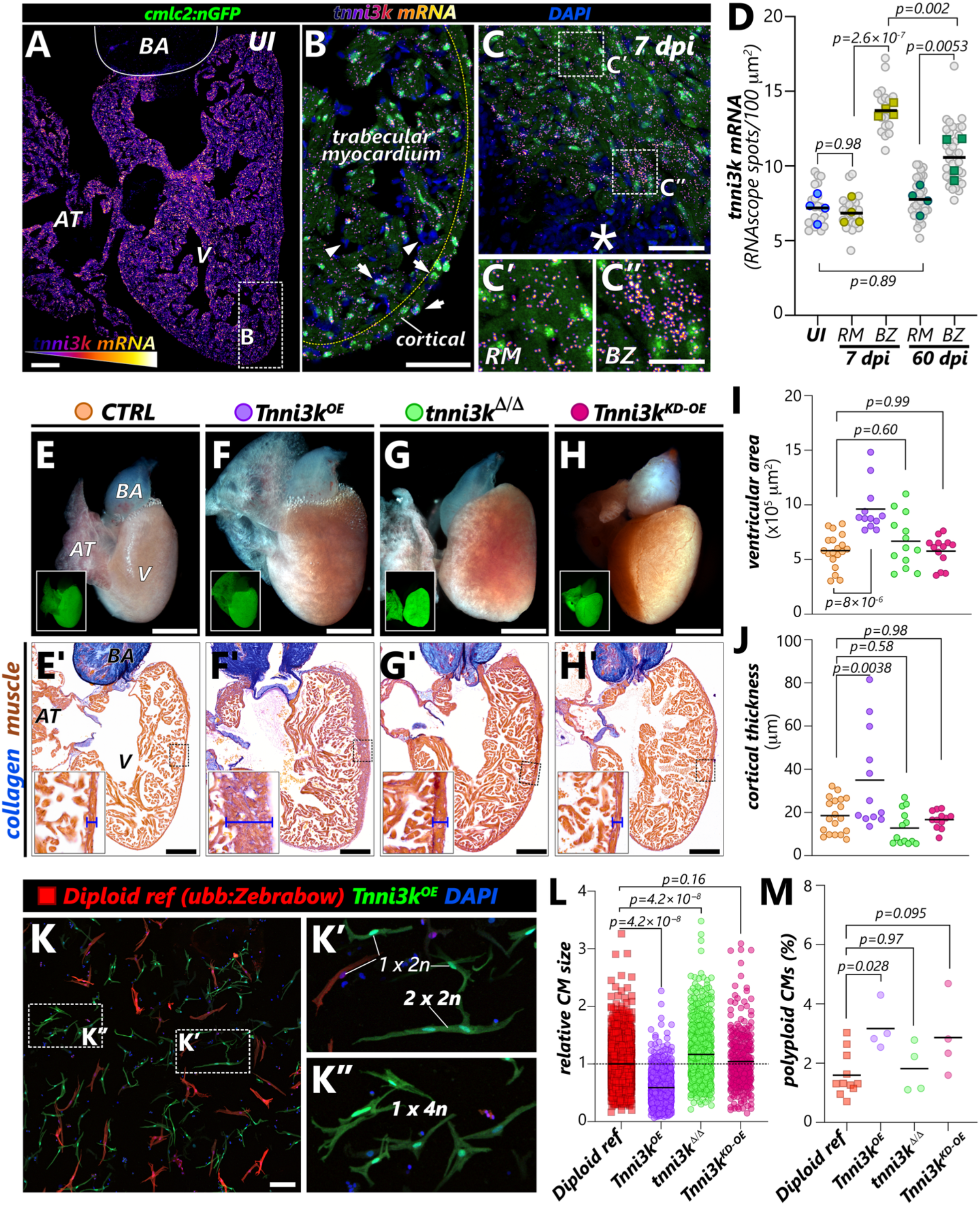
Elevated Tnni3k drives ventricular hyperplasia with only modest effects on cardiomyocyte ploidy in zebrafish. **(A)** Representative *tnni3k* RNA *in situ* hybridization (RNAscope) image of a wild-type zebrafish heart section. **(B)** Magnification of ventricle area boxed in (A). Arrows, cardiomyocyte nuclei; arrowheads, non-myocardial cells. **(C)** *tnni3k* RNAscope images of representative ventricular sections at 7 days after cryoinjury (dpi). Magnification of boxed areas is shown in C’ and C’’. Asterisk, injury area. **(D)** Quantification of *tnni3k* gene expression as RNAscope dots per area for uninjured (UI) and the indicated time points after injury. Each grey dot represents the mean for a given section (3 areas per section); colored dots represent the mean for individual animals (3-6 sections per animal, n =4). Black line, average. *p* values: one-way ANOVA, followed by Tukey’s multiple comparison test. **(E-H)** Representative whole heart images from zebrafish hearts of the indicated transgenic and mutant lines. GFP fluorescence from the transgene is shown in the bottom-left corner of the images. **(E’-H’)** Acid Fuchsin Orange G (AFOG) stained heart sections of the indicated cohorts. Magnifications of the boxed areas are shown in the bottom-left corner. **(I, J)** Quantification of ventricular area (I) and cortical thickness (G) on AFOG-stained tissue sections from the indicated cohorts. **(K)** Representative cardiomyocyte dissociation of a diploid reference from *ubb:Zebrabow* ventricles and *Tnni3k^O^*^E^ ventricles stained with the DNA dye DAPI for cardiomyocyte size and ploidy quantifications. Magnifications of the boxed areas are shown in (K’-K’’) to illustrate different ploidy classes. **(L, M)** Quantification of cardiomyocyte size (L) and frequency of polyploid cardiomyocytes (M) from ventricular dissociations of the indicated cohorts (3 pooled ventricles per sample, ∼1,000 CMs analyzed per sample). *p* values: One-way ANOVA, followed by Tukey’s multiple comparison test. AT, atrium; BA, bulbus arteriosus; BZ, border zone; CM, cardiomyocyte; RG, regenerated myocardium; RM, remote myocardium; V, ventricle. Scale bars: 100 μm (A, B and C), 25 μm (C’ and C’’), 1 mm, (E-H), 200 μm (E’-H’’), 100 μm (K).

We next tested whether *tnni3k* expression changes during zebrafish heart regeneration^22^. Given that elevated Tnni3k has been associated with cytokinesis failure, we hypothesized that the expression of this gene would be reduced in border zone (BZ) cardiomyocytes, which re-enter the cell cycle, divide, and give rise to new myocardium^1,2,23,24^. Unexpectedly, at 7 days post-cryoinjury (dpi), which corresponds to the peak of cardiomyocyte proliferation^22^, we detected a ∼2-fold increase in *tnni3k* expression in the BZ compared to the remote myocardium (RM) and uninjured controls **(Fig. 1C,D)**. By 60 dpi, an advanced stage of regeneration, *tnni3k* expression was partially reduced but did not return to basal levels **(Fig. 1D, S1C)**. In the remote myocardium, *tnni3k* expression did not differ significantly from that of uninjured controls. Thus, in a regenerative animal with diploid cardiomyocytes, *tnni3k* is naturally expressed across the myocardium, and its expression increases in proliferating cardiomyocytes, suggesting that Tnni3k’s role in ploidy regulation may depend on dosage or cellular context, rather than expression alone.

Because both elevated and reduced Tnni3k have been linked to cardiac pathology, we next tested whether changes in Tnni3k levels affect cardiac homeostasis and ploidy in zebrafish. To this end, we generated three new Tnni3k gain- and loss-of-function zebrafish models. First, we established a transgenic line expressing a bicistronic cassette encoding nuclear GFP and zebrafish *tnni3k* fused to six copies of the myc-tag under the cardiomyocyte-specific promoter *cmlc2* (*Tnni3k^OE^)* **(Fig. S1D, S1E)**. Second, because phenotypes described in murine models of Tnni3k overexpression depend on its kinase activity^17^, we engineered a kinase-dead transgenic line (*Tnni3k^KD-OE^*) carrying a K490R substitution in the canonical ATP-binding site^25^ **(Fig. S1D)**. Finally, we used CRISPR-Cas9 to delete the entire *tnni3k* locus to generate a mutant line (*tnni3k*^Δ/Δ^, **Fig. S1F**). RNAscope and qPCR confirmed a ∼30-fold increase of *tnni3k* in Tnni3k^OE^ and Tnni3k^KD-OE^ ventricles, and undetectable expression in *tnni3k*^Δ/Δ^ animals **(Fig. S1G-S1K)**.

All transgenic and mutant animals in this allelic series developed normally and exhibited standard survival rates. Whole-heart imaging and histological analysis revealed ventricular enlargement in Tnni3k^OE^ animals **(Fig. 1E,F,I)**, consistent with hypertrophic phenotypes reported in mice^26^. We also detected expansion of the cortical myocardium and collagen accumulation at the epicardium and between the cortical and primordial myocardium **(Fig. 1E’,F’,J)**. These features were not observed in *Tnni3k^KD-OE^* or *tnni3k*^Δ/Δ^ hearts **(Fig. 1G-H’,I,J).** To determine whether elevated Tnni3k promotes myocardial hypertrophy, we next measured cardiomyocyte area from ventricular dissociations across our allelic series. This analysis revealed an inverse correlation between Tnni3k levels and cardiomyocyte size. Compared with controls, cardiomyocytes were ∼41% smaller in Tnni3k^OE^ and ∼17% larger in *tnni3k*^Δ/Δ^ animals **(Fig. 1K,L)**. Cardiomyocytes from Tnni3k^KD-OE^ animals were indistinguishable in size from controls. Thus, larger ventricles built from smaller cardiomyocytes suggest that the overexpression of Tnni3k induces hyperplasia rather than hypertrophy in zebrafish.

Given that overexpression of murine Tnni3k has been reported to drive cardiomyocyte polyploidization in zebrafish^8^, we next tested whether changes in Tnni3k levels in zebrafish induce a shift in cardiomyocyte ploidy. To measure cardiomyocyte ploidy, we dissociated ventricles from our allelic series along with an internal diploid reference labeled with a red fluorescent protein^4^ **(Fig. 1K)**. We detected a small but significant increase in the polyploid fraction in Tnni3k^OE^ animals (∼1.7% in controls vs. ∼3.1% in Tnni3k^OE^, **Fig. 1M**). In contrast, ploidy in *tnni3k*^Δ/Δ^ and Tnni3k^KD-OE^ animals was indistinguishable from controls. This ∼2-fold increase in polyploid cardiomyocytes is markedly smaller than the ∼15-fold expansion previously reported in zebrafish overexpressing mouse Tnni3k^8^. It also falls well below the 45% threshold we previously described as affecting myocardial proliferation after injury^4^. Collectively, our data demonstrate that elevated Tnni3k induces several alterations in the uninjured zebrafish heart, including ventricular enlargement, cardiomyocyte hyperplasia, and fibrotic tissue accumulation, with only marginal changes in ploidy.

### Elevated Tnni3k drives fibrosis and prevents scar resolution after cryoinjury in a kinase-dependent manner

While we found that elevated *tnni3k* can be detected in regenerating cardiomyocytes and that forced expression produces myocardial hyperplasia, we also observed increased fibrosis in the Tnni3k^OE^ uninjured ventricles. These observations raised the question of whether perturbing Tnni3k levels would accelerate myocardial regeneration or impair it due to excessive scarring. To test this, we performed cryoinjuries in adult zebrafish from our Tnni3k allelic series, alongside controls, and analyzed their regenerative and fibrotic responses at different times post-injury.

As expected, control animals exhibited a transient fibrotic response that regressed at advanced stages after cryoinjury^22^. At 7 dpi, we detected the initial formation of the fibrotic scar, which peaked by 14 dpi **(Fig. 2A,E,F)**. By 60 dpi, control animals had largely resolved this transient scar, replacing it with regenerated myocardium **(Fig. 2A,G)**. As described^22,27^, only some residual collagen remains at this stage, intermingled with the thickened cortical myocardium. Tnni3k^OE^ animals showed a much more fibrotic response. At 7 dpi, fibrotic tissue accumulation was already ∼1.6-fold higher than in controls (5.57 ± 2.20% vs 3.36 ± 1.51%, mean ± SD; **Fig. 2B,E**). The difference became more pronounced by 14 dpi, when we detected a 2.3-fold increase in fibrotic tissue compared to controls (11.55 ± 2.04% vs 5.03 ± 2.08%, mean ± SD; **Fig. 2B,F**). By 60 dpi, Tnni3k^OE^ animals had failed to regenerate the myocardial wall and instead retained extensive scar tissue, ∼3.6-fold larger than in controls (4.25 ± 2.16% vs 1.19 ± 0.89%, mean ± SD; **Fig. 2B,G**). Thus, high levels of Tnni3k exacerbate the acute fibrotic response and impair scar regression, leading to failed ventricular wall regeneration. Consistently, we also documented increased fibrosis in mice overexpressing human *TNNI3K* in cardiomyocytes, both in the absence of injury and at 21 days after coronary artery ligation **(Fig. S2)**.

**Figure 2.**
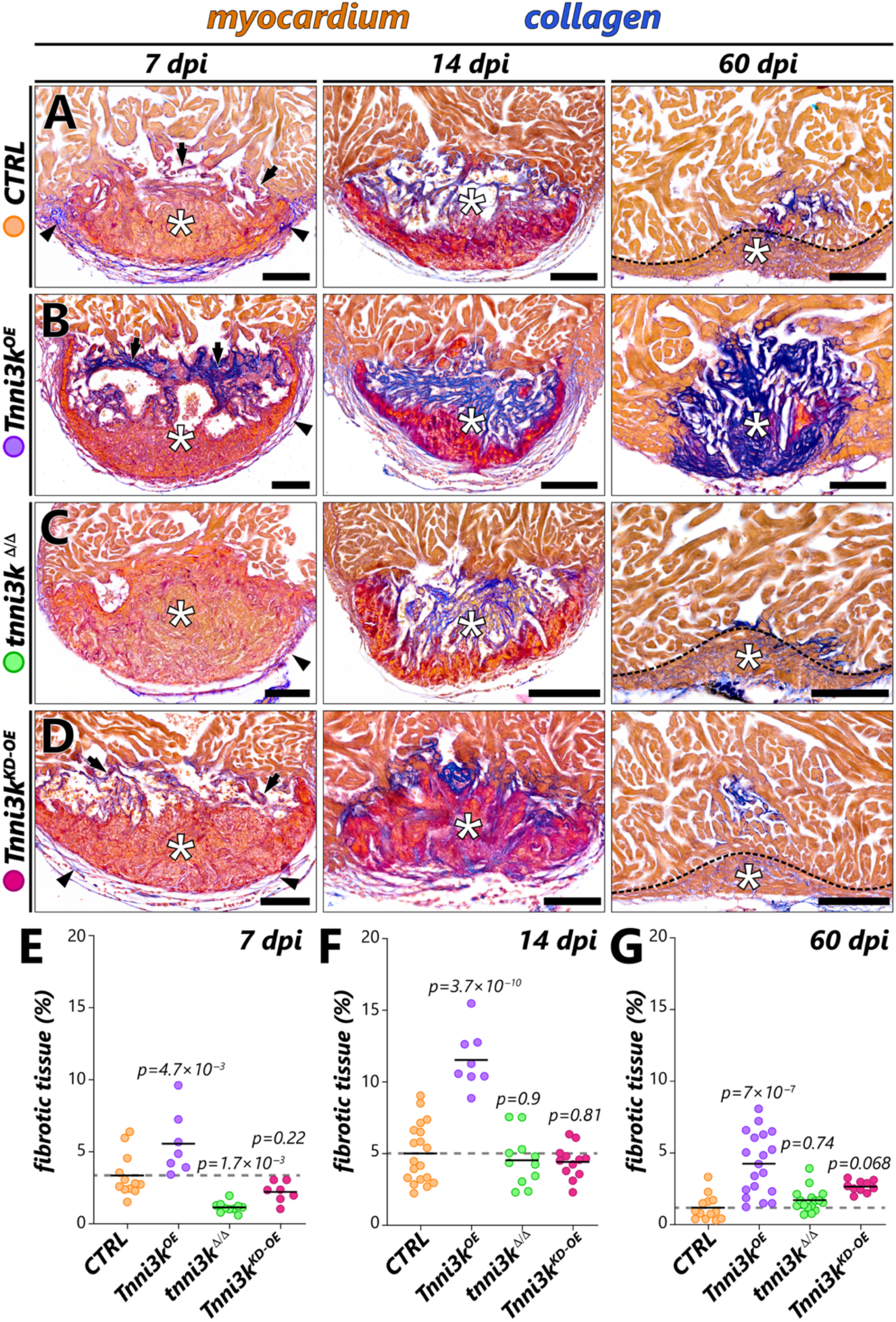
Elevated Tnni3k drives fibrosis and prevents scar resolution after cryoinjury in a kinase-dependent manner. **(A-D)** Representative heart sections from *CTRL* **(A)**, *Tnni3k^OE^* **(B)**, *tnni3k*^Δ/Δ^ **(C)**, and *Tnni3k^KD-OE^* **(D)** fish at 7, 14, and 60 days post-cryoinjury (dpi) stained with the Acid Fuchsin Orange G (AFOG) technique. Arrowheads, epicardial fibrosis. Asterisks, injured area. **(E-G)** Quantification of the fibrotic scar area, as the percentage of fibrotic tissue relative to ventricular area, in hearts from the indicated cohorts at 7-(E), 14-(F) and 60-(G) dpi. n(7 dpi) = 12, 9, 10, 7; n(14 dpi) = 19, 8, 11, 12; n(60 dpi) = 13, 11, 16, 10. *p-*values (relative to control): one-way ANOVA, followed by Tukey’s multiple comparison test. Scale bars, 100 μm.

In contrast to Tnni3k^OE^, both *tnni3k*^Δ/Δ^ and Tnni3k^KD-OE^ animals showed robust regeneration with complete recovery of the myocardial wall. At 7 dpi, *tnni3k*^Δ/Δ^ ventricles exhibited almost undetectable levels of scar, with ∼66% less fibrotic tissue than controls (1.71 ± 0.81% vs 3.36 ± 1.51%, mean ± SD; **Fig. 2C,E)**. Although this points to a role for Tnni3k in driving early fibrotic deposition, we found no differences in fibrosis by 14 dpi, and it remained indistinguishable from controls at 60 dpi **(Fig. 2C,F,G)**, demonstrating that loss of *tnni3k* does not impair regenerative capacity. Tnni3k^KD-OE^ animals followed the control trajectory across all three timepoints (**Fig. 2D-G**). Because Tnni3k^KD-OE^ animals express Tnni3k at levels matched to Tnni3k^OE^ **(Fig. S1K)** but lack kinase activity, these results indicate that the kinase activity is critical for the pro-fibrotic phenotype. Collectively, these data demonstrate that elevated Tnni3k blocks cardiac regeneration in the adult zebrafish in a kinase-dependent manner, and that loss of Tnni3k is tolerated without compromising regeneration.

### Regeneration failure in Tnni3k^OE^ animals cannot be explained by cardiomyocyte cell-cycle or ploidy defects

Given that our Tnni3k^OE^ animals failed to regenerate, we next sought to determine whether elevated Tnni3k impairs cardiomyocyte proliferation. A prior report attributed a comparable regeneration defect to polyploidization-driven loss of cardiomyocyte proliferation in zebrafish overexpressing mouse Tnni3k^8^. However, we previously demonstrated that ventricles with up to ∼45% polyploid cardiomyocytes regenerate robustly^4^. Because this is well above the modest ploidy increase in our Tnni3k^OE^ model **(Fig. 1M)**, we hypothesized that polyploidization alone could not account for the regeneration failure.

To test whether myocardial proliferation was affected, we examined cardiomyocyte cell-cycle activity across our allelic series at 3 and 7 dpi. At 3 dpi, co-staining for the cardiomyocyte-specific transcription factor Nkx2.5 and the proliferation marker PCNA revealed a ∼2-fold increase in BZ cardiomyocyte proliferation in *Tnni3k^OE^*animals, consistent with the enhanced S-phase entry reported in mice^19^ **(Fig. 3A,B)**. No changes were detected in *tnni3k*^Δ/Δ^ and Tnni3k^KD-OE^ animals. However, at 7 dpi, we detected equivalent percentages of cycling cardiomyocytes across all genotypes **(Fig. S3)**. Similarly, cardiomyocyte redifferentiation, measured by the re-expression of embryonic myosin isoforms shortly after proliferation completes^28^, was indistinguishable across the four genotypes **(Fig. 3C,D)**. Thus, differences in Tnni3k levels do not reduce myocardial proliferation or redifferentiation after injury. Instead, elevated Tnni3k accelerates cell-cycle entry shortly after injury.

**Figure 3.**
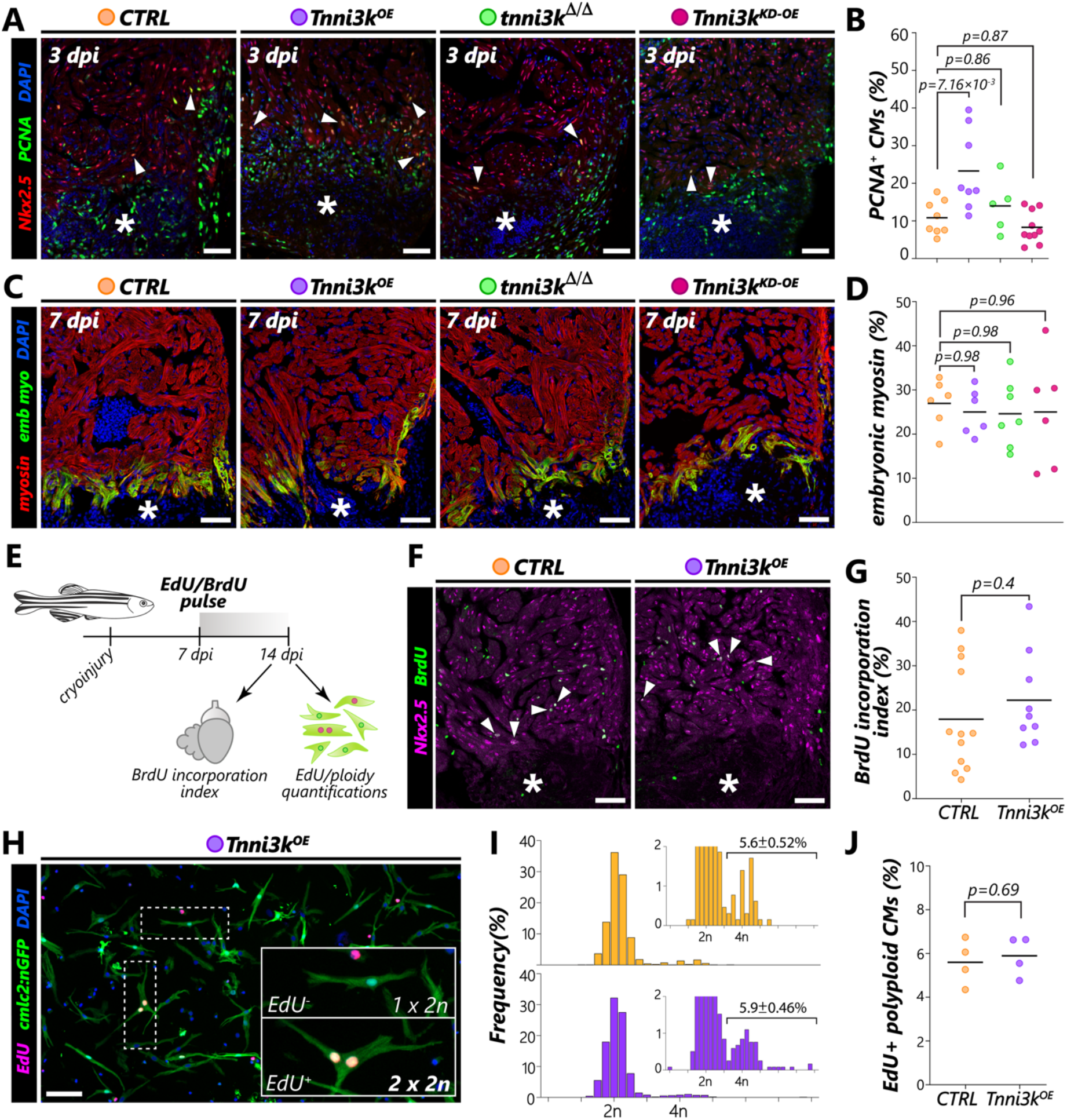
Regeneration failure in *Tnni3k^OE^* animals cannot be explained by cardiomyocyte cell-cycle or ploidy defects. **(A)** Heart sections from *CTRL*, *Tnni3k^OE^*, *tnni3k*^Δ/Δ^ and *Tnni3k^KD-OE^* fish at 3 days post-cryoinjury (dpi) immunostained for Nkx2.5 (cardiomyocyte nuclei) and PCNA (proliferating cell nuclear antigen, cycling cells). Arrowheads, Nkx2.5^+^ PCNA^+^ cardiomyocytes; asterisks, injury zone. **(B)** Quantification of cycling cardiomyocytes at 3 dpi as the percentage of cardiomyocytes positive for the PCNA marker (Nkx2.5^+^, PCNA^+^) relative to the total number of cardiomyocytes (Nkx2.5^+^) within a defined area (150 μm x 400 μm) in the injury border zone. *p* values: one-way ANOVA followed by Tukey’s multiple comparison test. 3 sections analyzed/sample. **(C)** Heart section images at 7 dpi immunostained to detect cardiomyocyte myosin heavy chain and the embryonic form of myosin heavy chain (MyHCIIa and third developmental stage MyHCI). Asterisks, injury zone. **(D)** Quantification of embryonic myosin heavy chain expressing myocardium relative to myocardium area within defined areas (200 μm x 400 μm) in the border zone of the injury at 7 dpi. *p* values: one-way ANOVA followed by Tukey’s multiple comparison test. 3 sections analyzed/sample. **(E)** Representation of the workflow for EdU/BrdU pulse and chase experiments. **(F)** Representative images of Nkx2.5 and BrdU immunofluorescence of *CTRL* and *Tnni3k^OE^* pulsed hearts at 14dpi. Arrowheads, Nkx2.5^+^ BrdU^+^ cardiomyocytes; asterisks, injury zone. **(G)** Percentage of BrdU^+^ cardiomyocytes, relative to the total number of cardiomyocytes (Nkx2.5^+^) in the border zone of the injury (150 μm x 400 μm areas) in *CTRL* and *Tnni3k^OE^* fish. *p* values: two-tailed, unpaired t-test. 3 sections analyzed/sample. **(H)** Representative image of *Tnni3k^OE^* fish ventricles dissociation at 14dpi (7 days after EdU pulse) stained with the DNA dye DAPI and to detect EdU. **(I)** Distribution of EdU^+^ cardiomyocytes DNA content in pulsed hearts at 14 dpi in *CTRL* and *Tnni3k^OE^* fish heart dissociations. Magnification to highlight low-frequency events is also shown. (n, *CTRL*= 643 EdU^+^ cardiomyocytes; n, *Tnni3k^OE^*= 1179 EdU^+^ cardiomyocytes). **(J)** Frequency of polyploid EdU^+^ cardiomyocytes in *CTRL* and *Tnni3k^OE^* heart dissociations. *P* values: two-tailed, unpaired t-test. 3 pooled ventricles per sample. Scale bars, 50μm (A, C, and F), 100μm (H).

PCNA labels cells undergoing DNA synthesis, but it does not distinguish whether these cells complete division or instead fail karyokinesis or cytokinesis. To track cardiomyocytes passing through S-phase, we pulsed them with the thymidine analogs BrdU or EdU at 7 dpi and analyzed them 7 days later **(Fig. 3E)**. Given that the Tnni3k^OE^ group was the only one to show a regeneration defect, we focused on this genotype, along with controls, for our experiments. In sections, we detected no differences in BrdU incorporation in BZ cardiomyocytes **(Fig. 3F,G)**, indicating that cardiomyocytes labeled during the pulse phase expanded comparably in both genotypes. Next, to determine whether cycling cardiomyocytes had completed cytokinesis or had instead become polyploid, we quantified ploidy in EdU+ cardiomyocytes from ventricular dissociations. Approximately 5% of EdU+ cardiomyocytes became polyploid after re-entering the cell cycle, but the vast majority of labeled cardiomyocytes efficiently divided and remained diploid in both groups **(Fig. 3H-J)**.

Our results using zebrafish Tnni3k overexpression contradict a previous report using mouse Tnni3k overexpression^8^. To test whether the mammalian and zebrafish Tnni3k orthologs function equivalently, we recreated a zebrafish line overexpressing mouse Tnni3k in cardiomyocytes (*m-Tnni3k^OE^*). This line reproduced all phenotypes we had characterized in our zebrafish overexpression line, including enlargement of the ventricles and expansion of the cortical layer **(Fig. S4A-S4E)**, cardiomyocyte hyperplasia, and a significant increase in the frequency of polyploid cardiomyocytes, but of the same modest magnitude as in our original line **(Fig. S4F-S4I)**. We also documented excessive collagen deposition at 7 dpi and failure to regress the scar at 60 dpi **(Fig S4J-S4K)** while maintaining proliferation and redifferentiation levels indistinguishable from controls **(Fig. S4L,S4M)**. These results demonstrate that both the zebrafish and mouse versions of Tnni3k induce equivalent phenotypes. Collectively, our results show that elevated Tnni3k does not reduce cardiomyocyte proliferation, redifferentiation, or productive cell division after injury. Both proteins increase cardiomyocyte ploidy only modestly, well below the ∼45% threshold compatible with robust regeneration^4^, demonstrating that the failure to regenerate in Tnni3k^OE^ animals cannot be attributed to a cell-cycle or ploidy defect.

### Elevated Tnni3k promotes basal inflammation, exacerbates the immune response after injury, and prevents its resolution

We next sought to determine what mechanisms underlie the regenerative failure induced by high levels of Tnni3k. Because the Tnni3k^OE^ phenotype is already apparent before injury, we hypothesized that a priming transcriptional state is already present at baseline. To test this, we performed RNA-sequencing of uninjured Tnni3k^OE^ and wild-type ventricles **(Fig. 4A)**. Differential-expression analysis identified 1,408 differentially expressed genes, of which 916 were upregulated and 492 downregulated in Tnni3k^OE^ **(Fig. 4B)**. *tnni3k* was among the most strongly induced transcripts in Tnni3k^OE^ hearts (log_2_FC=4.6, p-adj: 1.3×10⁻⁷⁰), confirming the experimental overexpression.

**Figure 4.**
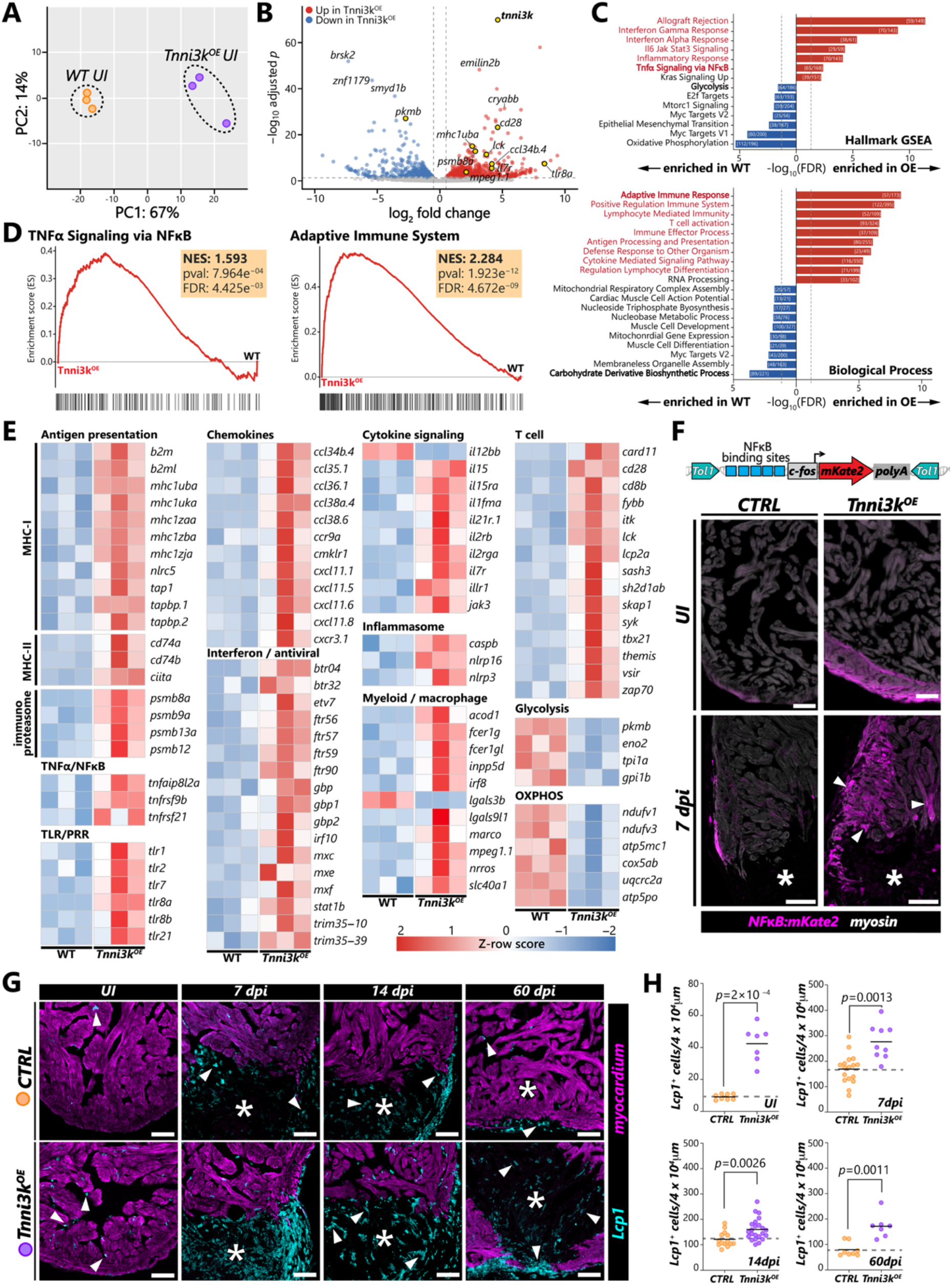
Elevated Tnni3k induces pro-inflammatory states in the uninjured heart and after injury. **(A)** Principal component analysis plot showing clustering of wild type and *Tnni3k^OE^* uninjured ventricles (3 samples/ experimental group; 3 pooled ventricles/sample). **(B)** Volcano plot showing differentially expressed genes (DEGs) between wild type and Tnni3k^OE^ ventricles. Blue and red dots represent statistically significant downregulated and upregulated genes in Tnni3k^OE^ samples, respectively. Selected genes are labeled. **(C)** Gene enrichment analysis (GSEA) bar charts showing enriched pathways in wild type versus Tnni3k^OE^ ventricles, using the Hallmark (top) and Gene Ontology Biological Process (bottom) gene sets. Red and blue bars denote pathways enriched in Tnni3k^OE^ or wild-type animals, respectively. Immune-related processes and pathways are highlighted in red. **(D)** Running enrichment-score curves for TNFα Signaling via NF-κB and the adaptive immune response as representative pathways upregulated in Tnni3k^OE^ ventricles. **(E)** Selected heatmaps representing relative expression of differentially expressed genes encoding immune system, inflammation, and metabolic related factors. Z-row scores of gene expression measurements are shown as colors ranging from blue to red. **(F)** NFκB activity reporter, carrying NFκB binding sites upstream of the minimal *c-fos* promoter, which drives the expression of mKate2. Immunofluorescence to detect NFκB activity in hearts of the indicated cohorts. Asterisks, injured area; arrowheads, NFκB-reporter+ myocardium. **(G)** Representative immunofluorescence images from *CTRL* and *Tnni3k^OE^* uninjured hearts and at 7, 14, and 60 days post-cryoinjury (dpi), detecting leukocytes (Lcp1^+^, arrowheads) and myocardium (myosin heavy chain). Asterisks, injured area. **(H)** Quantification of leukocyte number in hearts from the indicated cohorts and analyzed conditions. n (*UI)* = 8, 7; n (7 dpi) =17, 11; n (14 dpi) =14, 23; n (60 dpi) = 8, 7. *p* value*s*: two-tailed unpaired *t-test*. 3 sections analyzed/sample. Scale bars, 50 μm.

To identify key pathways differentially regulated in Tnni3k^OE^, we performed gene-set enrichment analysis **(Fig. 4C)**. Among the Hallmark gene sets, the most significantly upregulated terms in Tnni3k^OE^ ventricles were all immune-related. These included Interferon-γ and Interferon-α Responses, Inflammatory Response, and TNFα Signaling via NF-κB. Analysis of Gene Ontology Biological Processes identified antigen processing and presentation, adaptive immune response, lymphocyte-mediated immunity, and T-cell activation. Analysis of representative enrichment plots confirmed strong, coordinated enrichment across both arms of the immune system **(Fig. 4D)**. Conversely, glycolytic, oxidative, and contractile pathways were enriched in wild-type hearts, including oxidative phosphorylation, glycolysis, and muscle programs. Among the most strongly repressed transcripts was *pkmb* (log2FC=−2.73, p-adj: 8.3×10⁻²⁸; **Fig. 4B**), which encodes the zebrafish orthologue of the glycolytic enzyme pyruvate kinase M2 (PKM2).

Mining individual genes from our dataset, we found that essentially every major immune module was upregulated in the uninjured Tnni3k^OE^ heart **(Fig. 4E)**. These included the MHC class I and class II antigen-presentation machinery (*mhc1uba*, *b2m*, *tap1, cd74a/b*) and the immunoproteasome (*psmb8a*, *psmb9a*, *psmb13a*); an interferon/antiviral program (*mxe*, *mxc*, *stat1b*, *irf10*); Toll-like and pattern-recognition receptors (*tlr8a*, *tlr7*, *tlr2*); chemokines and their receptors (*ccl34b.4*, the *cxcl11* family, *ccr9a*); cytokine-signaling components (*il7r*, *il15*, *jak3*); a T-cell program (*cd28*, *lck*, *zap70*, *themis*, *tbx21*); and myeloid and macrophage markers (*mpeg1.1*, *marco*, *irf8*). We found a significant upregulation of components of the inflammasome, including *nlrp3, caspa,* and *caspbl*, which have previously been implicated in cardiomyocyte-induced inflammation and fibrosis^29^.

To test whether this inflammatory transcriptional signature reflected genuine pathway activity, we examined NF-κB activation *in situ* using *NFκB:mKate2* reporter. Reporter activity was elevated in uninjured Tnni3k^OE^ myocardium relative to controls and further amplified at 7 dpi **(Fig. 4F)**. This result corroborates the transcriptomic signature and localizes the pathway’s activity to the myocardium. Together, these data demonstrate that overexpression of Tnni3k in cardiomyocytes is sufficient to induce an inflammatory state in the absence of injury.

We next tested whether elevated Tnni3k increases the number of immune cells in the heart. In the absence of injury, leukocytes (identified using the pan-leukocyte marker Lcp1) were significantly more abundant (∼4-fold) in Tnni3k^OE^ ventricles than in controls **(Fig. 4G,H)**. These results demonstrate that high Tnni3k establishes a basal inflammatory state in the myocardium. Because both the initiation and resolution of inflammation are critical for zebrafish heart regeneration, we next tracked leukocyte infiltration dynamics after cryoinjury. Quantification of Lcp1⁺ cells across a region spanning the BZ myocardium revealed that Tnni3k^OE^ animals mounted an exacerbated inflammatory response, with significantly more leukocytes than controls at both 7 dpi (251.7 ± 82.97 vs 169.6 ± 57.5, mean ± SD) and 14 dpi (160.5 ± 42.36 vs 121.9 ± 29.85; **Fig. 4G-H**). Whereas the number of leukocytes in control animals continued to decrease at 60 dpi, the number of leukocytes was maintained at this time point in Tnni3k^OE^ hearts (172.1 ± 46.6 vs 79.7 ± 27.7; **Fig. 4G-H**). Thus, elevated Tnni3k promotes baseline inflammation, exacerbates it after injury, and impairs its resolution. This pattern mirrors the excessive, non-resolving fibrosis that these animals develop after injury.

In contrast to Tnni3k^OE^, neither *tnni3k*^Δ/Δ^ nor Tnni3k^KD-OE^ hearts displayed excessive inflammation, consistent with their normal fibrotic and regenerative outcomes. Leukocyte numbers were comparable to controls in both lines before injury and infiltration remained at or below control levels at 7 and 14 dpi **(Fig. S5A-S5F)**. By 60 dpi, leukocyte numbers had returned to control levels in both genotypes **(Fig. S5A, S5G)**. Thus, as with the fibrosis phenotype, the inflammatory phenotype is kinase-dependent. Importantly, we documented the same exacerbated inflammatory response in animals from the *m-Tnni3k^OE^* line, providing additional evidence that zebrafish and mouse Tnni3k behave indistinguishably **(Fig. S5A-S5G)**.

### Macrophage ablation before injury protects from Tnni3k-induced fibrosis and inflammation

Macrophages are essential for cardiac homeostasis and regeneration. In both zebrafish and mice, they also play an important role in the formation of the fibrotic scar^30^. Given the excessive fibrotic phenotype, we next tested whether elevated Tnni3k levels increase macrophage numbers in the heart. To quantify macrophage numbers, we performed *mpeg1.1* RNAscope *in situ* hybridization on sections of uninjured ventricles and hearts at 7 dpi, the peak of macrophage infiltration following cryoinjury^31^. Macrophage numbers were elevated in Tnni3k^OE^ hearts in both the uninjured state and at 7 dpi **(Fig. S5H-S5J)**, a phenotype we also observed in mice overexpressing *TNNI3K* **(Fig. S6)**. Our RNA-seq data also identified pro-inflammatory effectors within the upregulated immune signature, including the chemokine *ccl34b.4* **(Fig. 4B,E)**. In situ hybridization revealed more *ccl34b.4*+ immune cells and higher transcript levels in Tnni3k^OE^ hearts than in controls, both uninjured and at 7 dpi **(Fig. S5K-S5L)**. Together, these data establish a correlation between elevated Tnni3k, exacerbated inflammation, and the elevated fibrotic response to injury of the Tnni3k^OE^ model.

Macrophages deposit collagen directly into the scar in response to injury^30,31^. Given that we found increased macrophage numbers, we hypothesized that this increase is a key driver of the exacerbated, non-resolving fibrosis in Tnni3k^OE^ animals. To test this, we depleted macrophages with clodronate liposomes^32,33^ and analyzed fibrosis and leukocyte infiltration at 14 dpi, the peak of collagen accumulation in Tnni3k^OE^ hearts **(Fig. 2F)**. The initial regimen included two liposome injections before injury and one at 7 dpi (**Fig. 5A)**, which reduced both leukocyte numbers and fibrotic area at 14 dpi compared with PBS-treated Tnni3k^OE^ controls **(Fig. 5B-E)**. These results demonstrate that macrophages contribute to the exacerbated fibrotic phenotype of the Tnni3k^OE^ animals.

**Figure 5.**
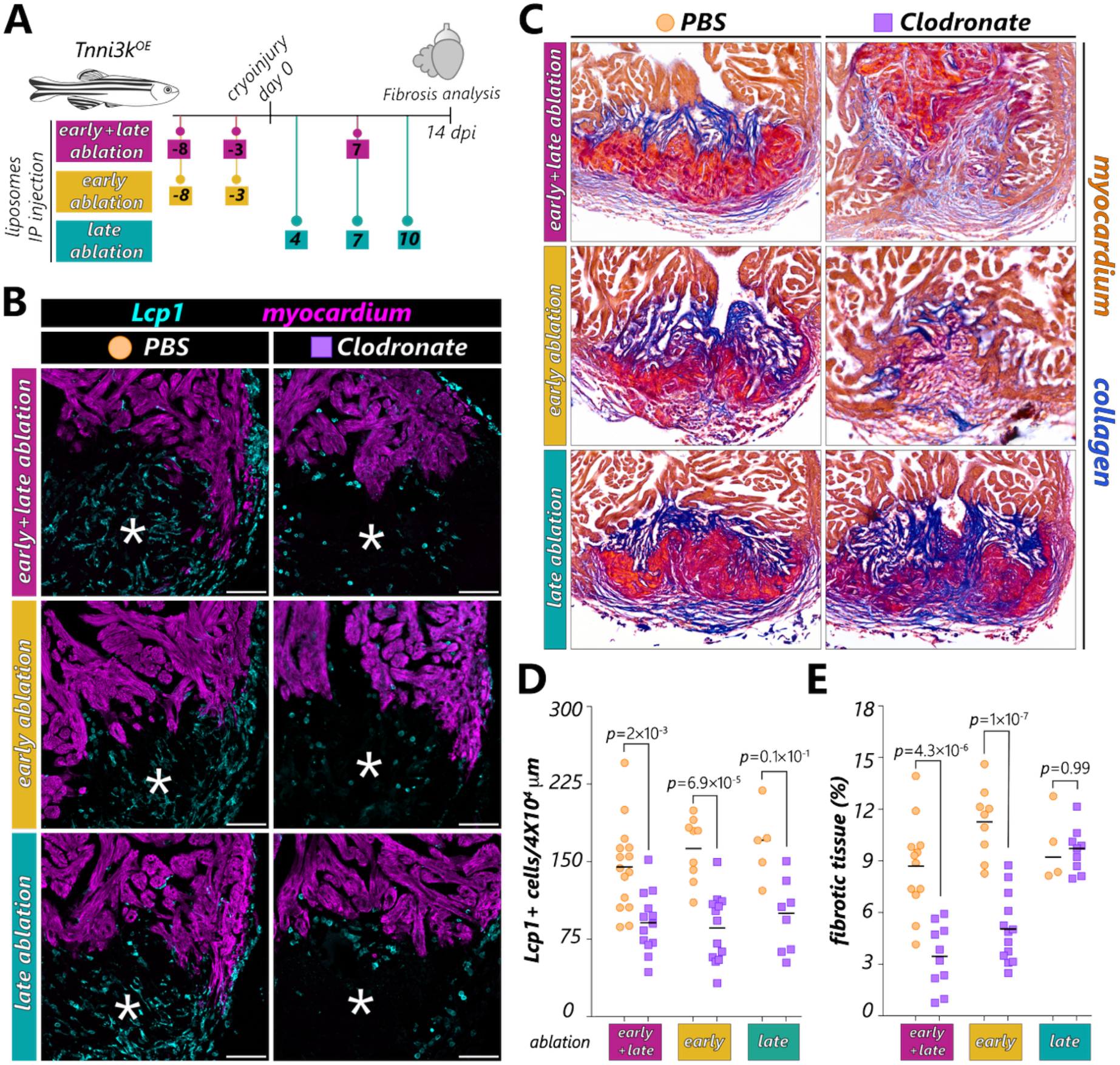
Macrophage ablation before injury protects from Tnni3k-induced fibrosis and inflammation. **(A)** Workflow showing the three tested regimens for macrophage ablation using liposomes on Tnni3k^OE^ zebrafish. **(B)** Leukocyte (Lcp1) and myocardium immunofluorescence images on Tnni3k^OE^ heart sections at 14 dpi. Asterisks, injury zone. **(C)** Representative AFOG-stained heart sections showing fibrotic tissue of the injury area at 14 dpi for the analyzed conditions. **(D)** Total Lcp1^+^ cell counts in 200 μm x 200 μm areas of the injury site. *p* values: one-way ANOVA, followed by Tukey’s multiple comparison test. 3 sections analyzed per sample. **(E)** Quantification of fibrosis after liposome injection at 14 dpi in Tnni3k^OE^ fish as the percentage of collagen-rich tissue area relative to ventricular area. *p* values: one-way ANOVA, followed by Tukey’s multiple comparison test. Scale bars, 100μm.

To dissect whether this aberrant fibrotic response was driven by the pre-injury inflammatory state or the post-injury macrophage influx, we designed two temporally distinct experimental regimens to ablate either early (pre-injury) or late (post-injury) macrophages **(Fig. 5A)**. While both regimens effectively reduced total leukocyte numbers at 14 dpi **(Fig. 5B,D)**, they had profoundly different impacts on fibrosis. The early ablation group showed a robust reduction in scar accumulation, while we found no differences in the late ablation group **(Fig. 5C,E).** The protective effect of early ablation points to the pre-injury macrophage pool as a critical determinant of the fibrotic response, consistent with the idea that a more pro-inflammatory environment enhances scarring^31^. These results identify the basal, Tnni3k-driven inflammatory state, rather than the post-injury macrophage response, as the primary driver of exacerbated fibrosis.

### Sustained Tnni3k overexpression after injury drives the fibrotic and inflammatory phenotype

We have established that elevated Tnni3k impairs zebrafish heart regeneration by promoting a pre-injury pro-inflammatory environment that amplifies after injury, resulting in exacerbated, non-resolving fibrosis. However, in all our experiments, animals continuously overexpressed Tnni3k before and after injury. Thus, we cannot determine whether the phenotype arises from the pre-injury state, from sustained overexpression during regeneration, or from both. To determine when Tnni3k must be elevated to drive fibrosis and inflammation, we next turned to tamoxifen-inducible Cre lines to switch overexpression “on” or “off” after cryoinjury.

In our original Tnni3k^OE^ line, the *nGFP-tnni3k* cassette is floxed and followed by an *mCherry* reporter **(Fig. S1D)**. When combined with a *Tg(cmlc2:CreER^T^*^2^) driver^2^ and treated with 4-hydroxytamoxifen (4-OHT), the *nGFP-tnni3k* cassette is removed, switching overexpression off (Tnni3k^OE^ inactivation). We also generated an activation transgene in which the two cassettes are swapped. The *cmlc2* promoter drives expression of a floxed *mCherry* cassette. Recombination leads to expression of the *nGFP-tnni3k* cassette, switching overexpression on. We treated animals with 4-OHT or vehicle at 5, 7, and 9 dpi, and analyzed fibrosis and inflammation at 21 dpi **(Fig. 6A)**.

**Figure 6.**
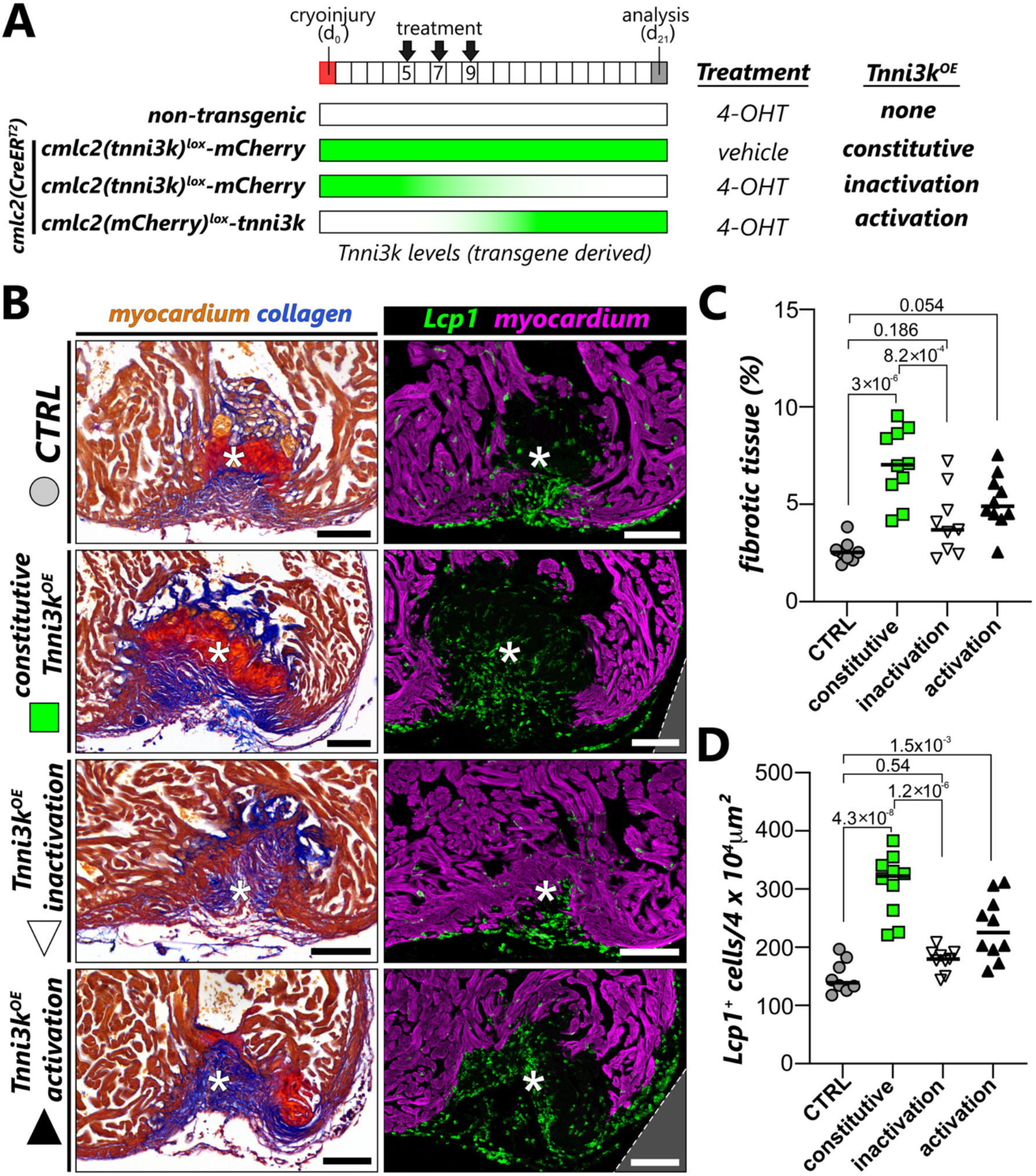
Sustained Tnni3k overexpression after injury drives the fibrotic and inflammatory phenotype. **(A)** DNA constructs and workflow used to modulate Tnni3k levels through tamoxifen inducible Cre recombination. **(B)** Representative AFOG stained heart sections (left column) and Lcp1 (pan-leukocyte marker), myosin heavy chain immunofluorescence images (right column) at 21 days post-injury (dpi). Asterisks, injury zone. **(C)** Quantification of the fibrotic scar area relative to total ventricular area in the analyzed conditions. *p* values: one-way ANOVA, followed by Tukey’s multiple comparison test. **(D)** Lcp1^+^ cell counts in defined areas (200 μm x 200 μm) of the injury site for all analyzed conditions. *p* values: one-way ANOVA, followed by Tukey’s multiple comparison test. 3 sections analyzed per sample. Scale bars, 100μm.

Consistent with our results at 7 and 14 dpi, constitutive Tnni3k^OE^ in animals treated with vehicle showed increased fibrosis at 21 dpi compared with non-transgenic controls (7.06 ± 1.84% vs 2.60 ± 0.59%, mean ± SD; **Fig. 6B,C**). Leukocyte infiltration was also doubled (307.2 ± 53.9 vs 150.0 ± 28.50; **Fig. 6B,D**), and the recovery of the myocardial wall was significantly impaired. We then asked whether continued overexpression is required to maintain this phenotype. Switching Tnni3k off after injury significantly reduced fibrosis towards control levels (4.10 ± 1.72%; **Fig. 6B,C**) and improved myocardial wall recovery. Leukocyte numbers were also lower in this condition (**Fig. 6B,D)**. Therefore, sustained Tnni3k overexpression after injury is necessary to maintain the exacerbated fibrotic and inflammatory responses.

We next tested whether raising Tnni3k levels post-injury is sufficient to recapitulate the phenotypes observed with constitutive overexpression. Switching overexpression on after cryoinjury increased both fibrosis and leukocyte infiltration above control levels, but these remained significantly lower than in the constitutive overexpression (5.16 ± 1.40% and 232.90 ± 53.79, respectively; **Fig. 6B-D**). Post-injury elevation is therefore sufficient to induce only a modest phenotype. Together with our macrophage ablation experiments, these results highlight the importance of the pre-injury basal inflammatory state as a key determinant of the Tnni3k^OE^-induced phenotype.

These results indicate that the development of the Tnni3k phenotype - exacerbated inflammation and fibrosis-has two requirements: inducing a basal inflammatory response and priming pre-injury macrophages, and maintaining high levels of Tnni3k after injury. Without pre-injury priming, no basal inflammation is induced, and the post-injury Tnni3k produces only a partial response. Switching Tnni3k off after injury normalizes fibrosis and inflammation, even in a heart that was primed before injury. The primed inflammatory state is therefore necessary but not sufficient; it drives exacerbated fibrosis only as long as Tnni3k signaling persists. Because switching off Tnni3k after injury permits the resolution of both fibrosis and inflammation, Tnni3k inhibition represents a candidate therapeutic strategy to improve outcomes following cardiac injury.

### Myocardial loss of *pkmb* phenocopies the Tnni3k^OE^ fibrotic and inflammatory phenotype

Our transcriptomic analysis identified *pkmb* as one of the most strongly repressed genes in Tnni3k^OE^ hearts **(Fig. 4B,E)**. Two features made *pkmb* a compelling candidate for linking elevated Tnni3k to the inflammatory phenotype. First, *pkmb* is the zebrafish orthologue of pyruvate kinase M2 (PKM2), a glycolytic enzyme with well-characterized functions beyond metabolism^34^. Second, PKM2 acts at the interface of metabolism and innate immunity. In endothelial cells, loss of PKM2 activates NF-κB signaling through RELB and, by depleting S-adenosylmethionine, lowers DNA methylation and de-represses endogenous retroviral elements. In turn, these responses trigger an interferon-driven antiviral response^35^. Because we found NF-κB signaling and an interferon/antiviral signature upregulated in Tnni3k^OE^ animals, we reasoned that *pkmb* repression could be a molecular link between elevated Tnni3k and myocardial inflammation. To test this, we generated a cardiomyocyte-specific *pkmb* mutant using the cardiodeleter system, a CRISPR/Cas9 approach restricted to cardiomyocytes recently pioneered by our laboratory^36^. A *pkmb*-guide shuttle carrying three guide RNAs, targeting exons 3, 8, and 11, was injected into *cmlc2:nGFP-Cas9* (*cardiodeleter*) animals **(Fig. 7A,B)**. Stable cardiomyocyte-specific mutants (*cardiodeleter*⁺ *pkmb-gs*⁺) developed normally to adulthood. RNAscope confirmed loss of *pkmb* mRNA relative to controls (cardiodeleter⁺/pkmb-gs⁻; **Fig. 7C**).

**Figure 7.**
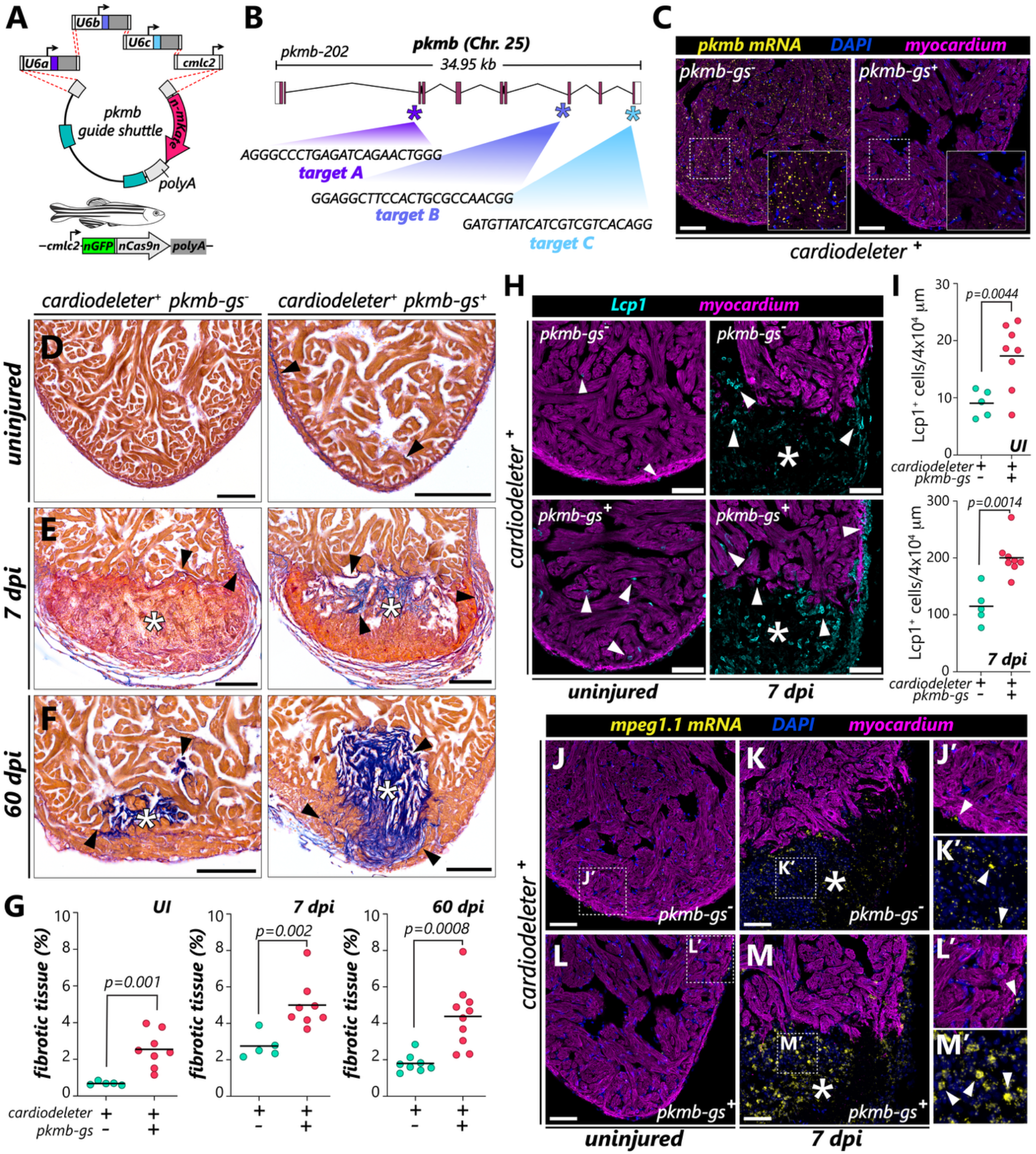
Myocardial loss of *pkmb* phenocopies the Tnni3k^OE^ fibrotic and inflammatory phenotype. **(A)** Components of the cardiodeleter system used to generate cardiomyocyte-specific *pkmb* mutants: the *pkmb* guide shuttle contains the three gRNAs downstream ubiquitous promoters U6a, U6b, and U6c, and the *cardiodeleter* zebrafish line, in which nuclear GFP and Cas9 expression is driven by the cardiomyocyte-specific promoter *cmlc2*. **(B)** Representation of the *pkmb* locus in the zebrafish genome. Filled boxes represent coding exons, empty boxes indicate the 5’ UTR and 3’ UTR. Asterisks indicate the location of the three selected gRNA targets. **(C)** *pkmb in situ* hybridization (RNAscope) on controls and cardiomyocyte-specific *pkmb* mutants in uninjured hearts. **(D-F)** Representative images of AFOG stained sections showing fibrosis (blue) in uninjured hearts (D), at 7dpi (E) and 60dpi (F) for the two analyzed cohorts. Asterisks, injured area. Arrowheads, fibrosis. **(G)** Quantification of fibrosis area relative to ventricular area. *p* value*s*: two-tailed unpaired *t-test*. **(H)** Immunofluorescence images showing leukocytes (Lcp1^+^) and myocardium of uninjured and 7 dpi hearts of the indicated cohorts. Asterisks, injury zone. Arrowheads, Lcp1^+^ cells. **(I)** Total Lcp1^+^ cell counts in 200 μm x 200 μm areas containing injury zone. *p* value*s*: two-tailed unpaired *t-test*. 3 sections analyzed per sample. **(J-M’)** Images of macrophage marker *mpeg1.1* RNAscope *in situ* hybridization co-detected with myosin heavy chain protein in uninjured heart and hearts at 7dpi. Magnifications of boxed areas are shown in J’-M’. Asterisks, injury zone, arrowheads, *mpeg1.1*^+^ cells. Scale bars, 50μm (C, G, and I), 100μm (D and E).

As in our Tnni3k^OE^, hearts lacking myocardial *pkmb* showed increased baseline fibrosis **(Fig. 7D,G)**. Moreover, these animals also showed more fibrosis after injury. At 7 dpi, fibrotic tissue in the injury zone was higher in *cardiodeleter*⁺ *pkmb-gs*⁺ animals than in controls **(Fig. 7E,G)**. Importantly, these hearts also failed to regenerate completely by 60 dpi and showed a ∼2-fold increase in scar retention **(Fig. 7F,G)**. The Tnni3k^OE^ inflammatory phenotype was also reproduced in these animals. Lcp1+ and *mpeg1.1+* cells were more abundant in *cardiodeleter*⁺ *pkmb-gs*⁺ hearts, both uninjured and at 7 dpi **(Fig. 7H-M)**. Loss of cardiomyocyte *pkmb* therefore reproduces the baseline and injury-induced features of the Tnni3k^OE^ heart: excess fibrosis, increased leukocytes, and macrophage accumulation. As with Tnni3k overexpression, *pkmb* deletion acts within cardiomyocytes yet drives the recruitment of non-myocardial immune cells. Our data identify *pkmb* repression as a candidate contributor to the fibrotic and inflammatory effects of high Tnni3k, and connect Tnni3k to an immunometabolic axis of cardiac injury.

## DISCUSSION

Although *Tnni3k* was discovered over 20 years ago, we know remarkably little about this cardiac-specific kinase, and many studies report conflicting results. Early studies showed that *Tnni3k* overexpression increased myogenesis, improved cardiac contractility *in vitro*^37^, and induced physiological hypertrophy *in vivo*^26^. In contrast, elevated Tnni3k has been found in samples from patients with dilated, hypertrophic, and ischemic cardiomyopathy^9–11^, and most studies in mammals demonstrate detrimental effects of elevated Tnni3k in response to various cardiac stresses^11,17,18^. In the context of regeneration, loss of this gene was initially associated with an increased proportion of diploid cardiomyocytes and higher levels of myocardial proliferation after injury^8^.

We initially approached this topic to determine the role of Tnni3k in a species with a natural capacity for regeneration. In adult mammals, elevated Tnni3k drives cardiomyocyte entry into S-phase but leads to polyploidization and worse outcomes after injury^19,20^. Our data demonstrate that in the highly regenerative zebrafish heart, elevated Tnni3k similarly accelerates cell-cycle entry, yet it drives productive division and hyperplasia rather than cellular hypertrophy or polyploidization. Despite a ∼30-fold increase in *tnni3k* expression, the proportion of polyploid cardiomyocytes remained well below the threshold required to impair regeneration^4^, and EdU tracing confirmed successful cytokinesis. We propose that this difference reflects the distinct cell-cycle competence of zebrafish and mammalian cardiomyocytes rather than a species-specific function of the kinase itself. If divisions are impaired, as in adult mammals, elevated Tnni3k may still favor S-phase entry but result in higher levels of polyploidization rather than bona fide divisions. This model also provides a mechanistic explanation for natural variation in mammalian heart ploidy: mouse strains with high Tnni3k expression may exhibit more polyploid cardiomyocytes simply because a higher proportion of their cells have entered the cell cycle and subsequently failed cytokinesis. Conversely, strains with low Tnni3k retain a diploid endowment because fewer cardiomyocytes are driven into S-phase. This interpretation differs from a previous report suggesting that mouse Tnni3k overexpression in zebrafish limits regeneration strictly by inducing polyploidy^8^. By re-evaluating both zebrafish and mouse orthologs, we ruled out ploidy and cell-cycle arrest as the primary drivers of regenerative failure. Instead, our findings pivot the mechanistic focus from a purely cell-autonomous cell-cycle defect to a dysregulation of the cardiac microenvironment.

Our analysis of Tnni3k^OE^ hearts identified a marked exacerbation of inflammation even in the absence of injury, and an increased fibrotic response that fails to regress. RNA-seq of uninjured Tnni3k^OE^ ventricles revealed a robust proinflammatory transcriptional program, including TNF/NF-κB signaling, the inflammasome, leukocyte activation, and immune-cell migration pathways. Consistent with this finding, Tnni3k^OE^ hearts accumulated more macrophages than controls under both homeostatic and post-injury conditions. The functional importance of this baseline inflammatory state was directly established by comparing two macrophage-depleting regimens: early ablation, targeting the pre-injury inflammatory environment, rescued the exacerbated fibrotic response in Tnni3k^OE^ animals, whereas late ablation initiated after injury had no effect. This temporal dissociation establishes that elevated Tnni3k creates a constitutive proinflammatory state that primes the heart for a maladaptive response. Using temporal genetic inactivation, we found that switching off Tnni3k overexpression after injury was sufficient to attenuate inflammation and fibrosis. Thus, while pre-injury priming sets the stage for aberrant healing, continuous Tnni3k signaling actively sustains the inflammation-fibrosis loop. Our data suggest that elevated Tnni3k levels in patients with cardiomyopathy may contribute to the inflammation observed in these subjects, and that inhibiting this kinase post-injury could halt this deleterious cascade.

The mechanistic link between elevated Tnni3k in cardiomyocytes and the activation of a non-cell autonomous inflammatory program remains an open question. However, inflammatory and fibrotic phenotypes are well described in the clinic and have been reproduced in animal models of cardiomyopathy affecting sarcomeric genes. Examples include models recreating mutations found in phospholamban^38^ (PLN^R9C/+^) and myosin heavy chains (MHC^R403Q^)^39^, which result in dilated and hypertrophic cardiomyopathy, respectively. As with Tnni3k, both genes are selectively expressed in cardiomyocytes, yet their most obvious effects are observed in non-myocardial cells. Profiling of PLN^R9C/+^ hearts revealed dysregulated metabolism in myocytes, followed by activation of innate and immune pathways in non-myocytes, and finally fibrosis and tissue stiffening^38^. Our transcriptomic data identified *pkmb*, a zebrafish orthologue of mammalian PKM, as one of the most consistently downregulated genes in Tnni3k^OE^ hearts. Cardiomyocyte-specific deletion of *pkmb* phenocopied the fibrotic and inflammatory features of Tnni3k^OE^ animals, identifying *pkmb* downregulation as a candidate effector downstream of elevated Tnni3k. Importantly, PKM2 has been implicated in coupling glycolytic metabolism to inflammatory gene expression in multiple cell types^35^, and metabolic reprogramming plays a central role in zebrafish heart regeneration^28^. Whether *pkmb* is the principal effector of Tnni3k-driven inflammation or one of several parallel mediators will require epistatic testing. Furthermore, identification of the substrates that connect Tnni3k kinase activity to NF-κB-target gene expression, inflammasome activation, and *pkmb* regulation is the next experimental priority for the field.

Beyond the specific role of Tnni3k in heart regeneration, what else can we learn from these results in the broader context of cardiac regeneration? Our results indicate that the basal inflammatory state is a critical determinant of regenerative outcome. Previous research has highlighted the immune system’s essential role in regeneration, yet we found that an exaggerated, chronic inflammatory response inhibits regeneration, even in the highly regenerative zebrafish. Identifying the molecular mechanisms by which cardiomyocytes induce this inflammatory state and how to interrupt the inflammation-fibrosis loop may define a critical therapeutic strategy for patients with heart failure. The systems established here may provide a platform to test the therapeutic potential of selective TNNI3K inhibitors for cardiac diseases driven by chronic myocardial inflammation.

## NON-STANDARD ABBREVIATIONS and ACRONYMS

4-OHT: 4-hydroxytamoxifen
BZ: border zone
cardiodeleter: cardiomyocyte-specific CRISPR/Cas9 deletion system (cmlc2:nGFP-Cas9)
cmlc2: cardiac myosin light chain 2 promoter
dpi: days post-cryoinjury
m-Tnni3kOE: mouse Tnni3k overexpression zebrafish line
*pkmb*: pyruvate kinase mb (zebrafish orthologue of PKM2)
*pkmb-gs*: pkmb-guide shuttle
RM: remote myocardium
Tnni3k: cardiac-specific troponin I-interacting kinase
Tnni3k^OE^: zebrafish Tnni3k overexpression transgenic line
Tnni3k^KD-OE^: kinase-dead Tnni3k overexpression transgenic line
*tnni3k*^Δ/Δ^: *tnni3k* CRISPR/Cas9 mutant line

## Acknowledgements

J.M.G.-R. thanks J. Butler, B. Howell, and the organizers of the 2026 Boston College Three-Day Sabbatical Retreat, for the opportunity to complete portions of this manuscript; the Boston College Animal Care Facility and past and present members of the González-Rosa Lab for fish care, preliminary data, and lab assistance; and Bret Judson of the Boston College Imaging Facility for training and assistance in image acquisition. Loren J. Field and Douglas A. Marchuk provided mouse sections for analysis.

## Sources of Funding

Work in the González-Rosa Lab is supported by the National Institutes of Health (R01HL164749), the American Heart Association (25IPA1453014), the Corrigan-Minehan Foundation (SPARK Award), the Hassenfeld Foundation (Hassenfeld Research Scholar), and internal funds from Boston College.

## Disclosures

The authors declare no competing interests.

## Supplemental Materials

Figures S1-S6

## MATERIALS AND METHODS

### Animals

Maintenance and growth of zebrafish (*Danio rerio*) embryos, larvae, and adults was carried out according to standard protocols approved by the Institutional Animal Care and Use Committees of Massachusetts General Hospital and Boston College. Ethical approval was obtained from the Institutional Animal Care and Use Committees of Massachusetts General Hospital and Boston College. For adult zebrafish experiments, animals ranging in age from 3 to 18 months, with approximately the same number of females and males, were used. Adult density was maintained at 3-4 fish·l^-1^ for all experiments in Aquarius racks, and fish were fed twice daily. Water temperature was maintained at 28 °C. Published strains used in this study include wild-type TuAB, *Tg(cmlc2:nGFP)^fb^*^18^ (ref. ^1^), *Tg(ubb:Zebrabow-M)^a^*^131^ (ref.^2^) and *Tg(cmlc2:nGFP-Cas9)^bcz^*^101^*^Tg^*(ref. ^3^). Details of the construction of the new lines generated in this study are described below. At least three independent founders of each line were isolated and tested to confirm the described expression patterns and phenotypes. All transgene sequences are available upon request.

### Generation of the Tnni3k^OE^ [Tg(*cmlc2:(nGFP-tnni3k)^lox^-mCherry)*] transgenic zebrafish line

To generate the Tnni3k^OE^ transgenic line, a construct containing the following DNA elements was assembled by Gibson cloning: (1) a 0.9 kb *cmlc2* promoter to drive specific expression in cardiomyocytes; (2) a floxed bicistronic *nlsGFP-P2A-6xmyc-tnni3k-polyA* cassette; and (3) an *mCherry-polyA* cassette. The transcript ENSDART00000129148.3 from *tnni3k* was amplified from cDNA extracted from adult TuAB zebrafish hearts using primers 5’-ATGGGGAATTACAAATCAA GACCTACTCAG-3’ and 5’-CTAGTTGCTGTCCTCAAAACT TCCATTG-3’ and cloned using the Zero Blunt TOPO PCR kit (K280002, Invitrogen) following the manufacturer’s instructions. To detect the presence of transgene-derived Tnni3k protein, six copies of the myc-tag were incorporated at the N-terminus of *tnni3k*. Tol2 flanking sites were used to maximize transgenesis. The resulting plasmid was sequenced and injected together with *Tol2* transposase mRNA in zebrafish embryos at the one-cell stage. In this line, all cardiomyocytes express a nuclear version of GFP and zebrafish Tnni3k. Upon Cre-mediated recombination, the floxed cassette is eliminated, and cardiomyocytes (and those cardiomyocytes that derive from them) are permanently labeled by the expression of mCherry. The official name of this line is *Tg(myl7:loxP-NLS-GFP-P2A-6xmyc- Dr.tnni3k-loxP-mCherry)^bcz^*^104^*^Tg^*.

### Generation of the Tnni3k^KD-OE^ [*Tg(cmlc2:nGFP-tnni3k^KD^)*] line

To generate the kinase-dead line, a construct containing the following DNA elements was assembled by Gibson cloning: (1) a 0.9 kb cmlc2 promoter to drive specific expression in cardiomyocytes; and (2) a bicistronic nlsGFP-P2A-6xmyc-tnni3k (K490R)-polyA cassette. The point mutation in the ATP-binding site was generated using the Q5 Site-Directed Mutagenesis Kit (E0554S, New England Biolabs). Tol2 flanking sites were used to maximize transgenesis. The resulting plasmid was sequenced and injected together with Tol2 transposase mRNA in zebrafish embryos at the one-cell stage. In this line, all cardiomyocytes express a nuclear version of GFP and a kinase-dead version of zebrafish Tnni3k. The official name of this line is *Tg(myl7:NLS-GFP-P2A-6xmyc-Dr.tnni3k(K490R))^bcz^*^105^*^Tg^*.

### Generation of the inducible Tnni3k^OE^ [Tg(*cmlc2:(mCherry)^lox^-nGFP-tnni3k)*] line

To generate the *cmlc2:(mCherry)^lox^-nGFP-tnni3k* transgenic line, we subcloned and reassembled in a different order the components of the construct *cmlc2:loxP-nls-GFP-P2A-6xmyc-Dr.tnni3k-loxP-mCherry,* described above. In this line, all cardiomyocytes express the red fluorescent protein mCherry. Upon Cre-mediated recombination, the *(mCherry)^lox^* cassette is eliminated, and cardiomyocytes (and those cardiomyocytes that derive from them) permanently express the fluorescent protein GFP directed to the nucleus and zebrafish Tnni3k. The official name of this line is *Tg(myl7:loxP-mCherry-loxP-NLS-GFP-P2A-6xmyc-Dr.tnni3k)^bcz^*^106^*^Tg^*.

### Generation and identification of the *tnni3k* locus deletion (Δ) allele

Guide RNAs (gRNAs) targeting 5’-AGGCGTCGTGATCCCTTGTCAGG-3’ (96 bp upstream of the first exon of *tnni3k* gene) and 5’-CCCATGAGCACCATGCGTTTCCA-3’ (37 bp upstream of the STOP codon) were identified using CRISPRScan^4^. gRNAs were produced as described^5^ using the specific primers 5’taatacgactcactataGGGCGTCGTGATCCCTTGTCgttttag agctagaa-3’, 5’-taatacgactcactataGGGAAACGCATGGTGCTCATgttttagagctagaa-3’ and the scaffold 5’-GATCCGCACCGACTCGGTGCCACTTTTTCAAGTTGATAACGGA CTAGCCTTATTTTAACTTGCTATTTCTAGCTCTAAAAC-3’. Both gRNAs (50 ng/μl each) and Cas9 protein (300 ng/μl, PNA Bio) were co-injected into one-cell-stage embryos as described^6^. For zebrafish genotyping, the Platinum II Hot-Start PCR Master Mix (Life Technologies) was used, following the manufacturer’s instructions. Primers flanking the guide RNA cut sites (forward: 5’-AGGAGTCTGATGATGCAGCTGTT-3’, reverse: 5’-TGACGTTCTTCAGAGATTTGCTGCT-3’) were used to identify allele deletion (Δ). These primers fail to produce an amplicon in wild-type animals because they are separated by ∼22 kb in the *tnni3k* endogenous locus but generate a ∼200 bp amplicon in animals carrying the Δ allele. To distinguish heterozygous and homozygous mutants, a second pair of primers (forward: 5’-TGCAGAGTCCCGCTTCCTGT-3’; reverse: 5’-GCGAAGGGAATCTCCCCCGT-3’) amplifying a region deleted in the Δ allele was designed. In this case, PCR generates a 435 bp amplicon only in animals carrying the wild-type allele. The official designation for the *tnni3k* Δ allele is *tnni3k^bcz^*^107^.

### Generation of the NFκB:mKate reporter

To detect cells with active NFκB-dependent gene transcription, a construct containing the following DNA elements was assembled by Gibson cloning: (1) six copies of the NFκB recognition sequence followed by the *c-fos* minimal promoter^7^ (a gift from John Rawls, Addgene plasmid # 44922); (2) the CDS of the red fluorescence protein mKate2 followed by a polyadenylation signal; and (3) a *crybb:mCherry-polyA* cassette as transgenesis control. *Tol1* flanking sites were used to maximize transgenesis. The resulting plasmid was sequenced and injected together with *Tol1* transposase mRNA in zebrafish embryos at the one-cell stage. In this line, cells in which the NFκB pathway is active express mKate2. All independent insertions for this construct reproduce the reported expression pattern, but a single insertion was used for all experiments. The official name of this line is *Tg(NFkB:mKate2; crybb:mCherry)^bcz^*^108^*^Tg^*.

### Construction of *pkmb guide shuttle (pTol1-U6abc-pkmb; cmlc2-nmKate-SV40pA)* and generation of the cardiomyocyte-specific *pkmb* mutant line

A *guide shuttle* encoding three gRNAs targeting zebrafish *pkmb* gene was generated as previously described^3^. Briefly, a construct containing the following DNA elements was assembled by HiFi cloning (NEB): (1) the linearized *Tol1-cm:n-mKate guide shuttle* backbone vector; (2) a *U6a:sgRNA* (*pkmb* A) cassette targeting the sequence AGGGCCCTGAGATCAGAACTGGG; (3) a U6b:sgRNA (pkmb B) cassette targeting GGAGGCTTCCACTGCGCCAACGG; and (4) a U6c:sgRNA (*pkmb* C) cassette targeting GATGTTATCATCGTCGTCACAGG. The resulting plasmid was sequenced and injected together with *Tol1* transposase mRNA in *cardiodeleter+* embryos at the one-cell stage. The F0 mosaics positive for both transgenes were screened, and at least three identified founders were selected. The resulting double positive F_1_ constituted the cardiomyocyte-specific *pkmb* mutant line used for further analysis. The official name of this line is *Tg(rnu6-32:CRISPR4-pkmb,rnu6-14:CRISPR5-pkmb,rnu6-7:CRISPR6-pkmb,myl7:NLS-mKate2)^bcz^*^110^*^Tg^*.

### Liposome intraperitoneal injection

Liposome injections were performed as described^8–10^. Fish were anesthetized with tricaine (0.032%; wt/vol; Sigma) and placed ventral side up on a foam sponge. Intraperitoneal (IP) injection of 10 μl of liposomes containing PBS or clodronate (5 mg/ml) (C-005, Liposoma) was performed using a syringe with ultra-fine needle (Beckton Dickinson). In all cases, hearts were dissected at 14 dpi and processed for histological analysis.

### Tamoxifen treatment in adult zebrafish

To induce Cre-ER^T2^-mediated recombination in *Tg(cmlc2:(nGFP-tnni3k)^lox^-mCherry)* and *Tg(cmlc2:(mCherry)^lox^-nGFP-tnni3k*) adult zebrafish, 5 μl of 200 μM 4-hydroxy-tamoxifen (4-OHT, Sigma) was administered orally at 5, 7, and 9 dpi. A 10 mM 4-OHT stock solution was preheated for 10 min at 65 ^°^C prior to preparing the working solution. Hearts were dissected and processed for analysis at 21 dpi.

### Adult zebrafish cardiac injuries

Cryoinjury was performed on adult zebrafish hearts as previously described^11^. Fish were anesthetized in tricaine (0.032%; w/v, Sigma), placed with their ventral side up on a sponge, and then a small incision was created to expose the apex of the ventricle. A Kimwipe was applied to the ventricle surface to remove excess water. Next, a platinum filament, previously cooled in liquid nitrogen, was placed on the ventricular surface and maintained for a few seconds until thawing to induce rapid freezing of approximately 20% of the ventricle. After this, animals were returned to a fish tank containing clean, fresh water for recovery. Resection was performed as described^12^. Following anesthesia and exposure of the ventricle as described above, ∼20% of the ventricle was removed using scissors. To prevent excessive bleeding, a Kimwipe was used to blot the incision immediately after the apex was removed. Following the injury, the fish were returned to fresh water.

### Zebrafish heart dissection and processing for histology

Adult zebrafish were euthanized by immersion in 0.16% tricaine (Sigma) and hearts dissected as described^11^. In brief, zebrafish were put ventral side up on a sponge submerged in ice-cold heparin-containing phosphate-buffered saline (PBS). After a deep cut into the aorta to promote bleeding, the entire heart was removed from the pericardial chamber and placed in a 0.5 M potassium chloride (KCl) solution in PBS for a few minutes. Whole heart samples were fixed by overnight incubation in 4% paraformaldehyde (PFA) in PBS at 4 °C. After fixation, samples were embedded in paraffin following conventional histological procedures, and 7μm paraffin sections were obtained using a semi-automated rotatory microtome (Leica) on Superfrost Plus slides (1255015, ThermoFisher Scientific) and kept at 4 °C until processed for histological staining.

### Fluorescent *in situ* hybridization on paraffin-embedded tissue sections

Codetection of gene transcripts and proteins in 7 µm paraffin-embedded heart sections was performed as described^3^ using RNAscope^TM^ (Advanced Cell Diagnostics, Biotechne). The Multiplex Fluorescent Reagent Kit v2 was used following the manufacturer’s instructions. The probes used were *Dr-tnni3k* (NM_001204823.1, *Danio rerio*, #805719), *Dr-mpeg1.1* (NM_212737.1, *Danio rerio*, #536171), *Dr-ccl34b.4* (NM_001115049.2, *Danio rerio*, #820541, *Dr-pkmb* (NM_001003488.1, *Danio rerio*, #1463751). In all cases, nuclei were counterstained with DAPI, and slides were mounted in FluorSave (Millipore). Images were obtained using Nikon A1, Zeiss LSM 880, or Leica Stellaris 5 confocal microscopes for further analysis.

### Total RNA isolation from zebrafish heart ventricles

Adult zebrafish hearts were dissected in ice-cold PBS and transferred into a small petri dish containing fresh ice-cold PBS. We used whole hearts for RT-qPCR, and isolated ventricles for RNA-seq. Samples were washed twice with ice-cold PBS. Sample pools, consisting of 2 hearts or 3 ventricles each, were then transferred to tissue homogenizing tubes (Bertin Corp. #P000911-LYSK0-A, CK28) containing 700 µl of QIAzol^TM^ Lysis Reagent (Qiagen) and lysed on a TissueLyser II (Qiagen) at 23 Hz for 10 min at 4°C. Lysates were stored at -80°C. For RNA extraction, nucleic acids were first recovered with MaXtract High Density (Qiagen, #129056) and total RNA was isolated by column purification with on-column DNA digestion (Qiagen, miRNeasy mini #1038703) according to the manufacturer’s instructions. RNA was eluted in 30 µL RNase-free water, and concentration was measured using a Nanodrop One (Thermo Scientific).

### Real time-quantitative PCR (RT-qPCR)

For gene expression analysis, total RNA was isolated from adult zebrafish hearts as described above. RNA was retrotranscribed to cDNA using the SuperScript™ IV VILO™ Master Mix (Invitrogen, #11756050). Real-time PCR was performed in a StepOnePlus™ real-time PCR system (Applied Biosystems) using Power SYBR Green PCR Master Mix (Applied Biosystems, Life Technologies)/TB Green® Premix Ex Taq™ (Perfect Real Time, Takara, #RR420A). Reaction mixtures were incubated for 30 seconds (sec) at 95°C, followed by 40 cycles of 5 sec at 95°C, 20 sec at 60°C, and finally a melting curve protocol. *tnni3k* expression was normalized to the ribosomal protein S11 (*rps11*) mRNA content using the comparative Ct method (2-ΔCt). In all cases, each PCR was performed with technical triplicates per sample. Each biological replicate consisted of a pool of 2 whole hearts, and at least 4 biological replicates were analyzed per genotype. *tnni3k* mRNA was detected using the primers 5’-GAGTTGGAATACGCCCTCAA-3’ and 5’-CATGTTCCTCCTCATCTCATCC-3’; rps11 was detected using the primers 5’-AGAAACAGCCCACCATCTTC-3’ and 5’-AGCCCAATCCAACGTTTCT-3’.

### Immunofluorescence on paraffin-embedded tissue sections

Immunofluorescence was performed as described previously^13^. In short, sections were deparaffinized and rehydrated, and antigen retrieval was performed by pressure-cooking in a citrate-based unmasking solution (H3300, Vector). Sections were incubated for 15 min in methanol, washed in 0.1% Tween-20 in PBS (PBSTw) and photobleached for 45 min. After washing with PBSTw, sections were permeabilized in 0.25% Triton X-100 in PBS (PBSTx) for 10 min. To minimize nonspecific antibody binding, sections were blocked in 5% goat serum and 5% bovine serum albumin in PBSTw for 1 h at room temperature. Sections were incubated with primary antibodies in blocking buffer overnight at 4°C. Primary antibodies used were mouse anti-tropomyosin (clone CH1, Developmental Studies Hybridoma Bank; 1:50), chicken anti-GFP (AVES, 1:500), mouse anti-GFP (clone B-2, Santa Cruz Biotechnology; 1:200), mouse anti-PCNA (clone PC10, Santa Cruz Biotechnology, 1:500), rabbit anti-nkx2.5 (GTX128357, GeneTex, 1:500), rabbit anti-ERG (ab110639, Abcam, 1:150), mouse anti-myosin heavy chain (clone MF20, Invitrogen, 1:100) rabbit anti-Lcp1 (GTX124420, GeneTex; 1:500), mouse anti-embryonic myosin (clone N2.261, 1:25), and rabbit anti-F4/80 (#30325, Cell Signaling, 1:400). Slides were washed in PBSTw before incubation with Alexa-conjugated secondary antibodies (Life Technologies, 1:500) in blocking buffer for 1 h at room temperature. In all cases, nuclei were counterstained with DAPI (Invitrogen), and slides were mounted in FluorSave (Millipore). Zeiss LSM 880 and Leica Stellaris 5 confocal microscopes were used for image acquisition.

### Acid fuchsin-orange G (AFOG) staining

Acid fuchsin-orange G (AFOG) stain was used to detect fibrotic tissue, as described^1^. In short, after deparaffination and rehydration, tissue sections were fixed in Bouin’s Fluid (Fisher Scientific) overnight at 60 ^°^C, followed by a wash of 5 min in 1% phosphomolybdic acid (Sigma). Then, sections were incubated in a preheated AFOG solution (5 g/l Water Blue, 10 g/l Orange G, and 15 g/l Acid Fuchsin in acidified water) for 15 min, dehydrated, and mounted in DPX mounting media (Sigma). Muscle, fibrin/cell debris, and collagen were stained brown, orange, red, and blue, respectively. Images of evenly spaced AFOG-stained serial sections of the whole heart were captured on a SLIDEVIEW™ VS200 (Olympus) using 40X and 60X lenses.

### Heart morphology analysis and fibrosis quantification

Morphological analysis of uninjured adult zebrafish hearts was performed on AFOG-stained sections. For ventricle area measurement, masks encompassing the entire ventricle were manually generated in Adobe Photoshop (Adobe Systems Incorporated), and the area was measured in ImageJ. For this quantification, an average of 10 sections per heart was analyzed (min=7, max=18). The thickness of the myocardial cortical layer was quantified in three sections per sample, with three points measured in each section. To quantify fibrosis area in regenerating hearts, an average of 10 AFOG-stained sections per heart (min=7, max=18) was analyzed. Masks containing the entire ventricular area and the scar area were manually generated using Adobe Photoshop (Adobe Systems Incorporated) based on differential staining (uninjured/regenerated muscle = brown/orange; fibrotic area = (fibrin (red) + collagen (blue)). Selected areas were measured using ImageJ software. The fibrotic area was normalized to the total ventricular area to calculate the percentage of the scar size for each heart.

### Cardiomyocyte Isolation from Zebrafish Ventricles

Individual cardiomyocyte suspensions were obtained as previously described^1^. Dissected ventricles were pre-digested on ice for 15 min in 0.2% trypsin, 0.8 mM EDTA (25200-056, Gibco) supplemented with 20 mM glucose and 10 mM 2,3-butanedione monoxime (BDM, B0753, Sigma) with gentle agitation. Then, the samples were digested for 45 min at room temperature in Accumax (SCR006, EMD Millipore) supplemented with 20 mM glucose and 10 mM BDM under mild agitation. Tissue fragments were dissociated by gentle pipetting, and cell suspensions were fixed in 10% neutral buffered formalin (HT501128, Sigma) for 1 hour at room temperature. Cells were pelleted by centrifugation at 400 x *g* for 5 minutes, resuspended in PBS, spread onto Superfrost Plus slides (1255015, Thermo Fisher Scientific), and air-dried. In all experiments, 3 ventricles were pooled per biological replicate. As the diploid reference, age-matched *ubb:Zebrabow* or wild-type ventricles were pooled and dissociated together with experimental samples.

### Cardiomyocyte size and DNA content quantification in Cell Spreads

Cardiomyocyte areas were measured in cell spreads obtained from ventricle dissociations. Binary masks containing cardiomyocyte areas were manually generated in Adobe Photoshop (Adobe Systems Incorporated). Damaged cells were excluded from the analysis. Independent binary masks were generated based on fluorescent protein expression to distinguish control and experimental cell populations. Masks were quantified using Fiji/ImageJ software. The area of experimental cardiomyocytes was normalized to the average area of the reference population. On average, 200-400 cardiomyocytes from the reference population and 600-700 from the experimental samples were measured. Cardiomyocyte DNA content was determined by quantifying the integrated nuclear density of cells stained with the DNA dye 4’,6-diamidino-2-phenylindole (DAPI), as described ^1^. For each biological replicate, tiled images were captured using a Nikon A1 or a Leica Stellaris 5 confocal microscope, and ∼1,000 experimental cardiomyocytes were quantified. Nuclear masks were manually generated in Adobe Photoshop (Adobe Systems Incorporated). Integrated density was calculated in ImageJ on a per-nucleus basis, and ploidy was estimated per cell. Isolated nuclei and damaged cells were excluded from the analysis. Independent binary masks were generated based on fluorescent protein expression to distinguish the experimental and reference populations, or based on the presence of EdU signal (see below).

### Bromodeoxyuridine and Ethynyldeoxyuridine pulse-chase experiments

For 5-bromo-2-deoxyuridine (BrdU) pulse-chase experiments, adult zebrafish were subjected to cryoinjury. At 7 days post-injury (dpi), animals were injected intraperitoneally with 50 μl of 2.5 mg/ml BrdU (B5002-1G, Sigma) in PBS. Hearts were dissected and processed for histologic analysis at 14 dpi. In the case of 5-Ethynyl-2′-deoxyuridine (EdU) pulse and chase experiments, adult zebrafish were intraperitoneally injected with 20 μl of EdU 10 mM in PBS at 7 dpi. At 14 dpi, cardiomyocyte dissociations were obtained as described above. Cardiomyocyte spreads were rehydrated in PBS and permeabilized for 45 min in 0.1% IGEPAL CA-630 (I8896, Sigma), 3% BSA in PBS. For EdU detection, the Click-iT Plus^TM^ reaction was performed according to the manufacturer’s instructions, and then the slides were stained for 30 min with DAPI dihydrochloride (Invitrogen) in permeabilization solution.

### Quantification of cardiomyocyte cell cycle entry and proliferation

The cardiomyocyte cell-cycle entry index was calculated in adult zebrafish hearts at 7 dpi from sections immunostained with anti-nkx2.5 (GeneTex) to label cardiomyocyte nuclei and anti-PCNA (Santa Cruz) to identify cycling cells. Nuclei were counterstained with DAPI. For each heart, three ventricular sections containing the largest injury areas were imaged and quantified. Positive cells for both markers (nkx2.5^+^, PCNA^+^), as well as total number of cardiomyocytes (nkx2.5^+^ cells) were counted manually using Fiji/ImageJ software in defined regions (150 μm x 400 μm) in the border zone of the injury. The percentages of nxk2.5^+^PCNA^+^/nxk2.5^+^ cells from individual sections were averaged to establish an index for each animal.

To calculate the BrdU labeling index, the same method was applied to 14 dpi ventricular sections from fish that received a BrdU pulse at 7dpi. Sections were immunostained with anti-nkx2.5 and anti-BrdU antibodies to detect Nkx2.5^+^ single-positive cardiomyocytes and Nkx2.5^+^, BrdU^+^ double-positive cardiomyocytes.

### RNA-seq

RNA quality assessment, library preparations, and sequencing reactions were conducted at GENEWIZ, LLC. (South Plainfield, NJ, USA). RNA samples were quantified using Qubit 2.0 Fluorometer (Life Technologies) and RNA integrity was checked using Agilent TapeStation 4200 (Agilent Technologies). RNA sequencing libraries were prepared using the NEBNext Ultra RNA Library Prep Kit for Illumina following the manufacturer’s instructions (NEB). Briefly, mRNAs were first enriched with Oligo(dT) beads. Enriched mRNAs were fragmented for 15 minutes at 94 °C. First strand and second strand cDNAs were subsequently synthesized. cDNA fragments were end-repaired and adenylated at 3’ ends, and universal adapters were ligated to cDNA fragments, followed by index addition and library enrichment by limited-cycle PCR. The sequencing libraries were validated on the Agilent TapeStation (Agilent Technologies) and quantified using a Qubit 2.0 Fluorometer (Invitrogen) and quantitative PCR (KAPA Biosystems). The sequencing libraries were clustered on 1 lane of a flowcell. After clustering, the flowcell was loaded on the Illumina HiSeq instrument (4000 or equivalent) according to the manufacturer’s instructions. The samples were sequenced using a 2×150 bp paired-end (PE) configuration. Image analysis and base calling were conducted by the HiSeq Control Software (HCS). Raw sequence data (.bcl files) generated by the Illumina HiSeq were converted to fastq files and demultiplexed using Illumina’s bcl2fastq 2.17 software. One mismatch was allowed for index sequence identification. On average, 20M reads were obtained per sample.

### Sequence mapping and Differential expression analysis

Sequence reads were trimmed to remove possible adapter sequences and low-quality nucleotides using Trimmomatic v.0.36. The trimmed reads were mapped to the *Danio rerio* GRCz10.89 reference genome available on ENSEMBL using the STAR aligner v.2.5.2b. Unique gene hit counts were calculated by using featureCounts from the Subread package v.1.5.2. DESeq2 package was used to normalize the counts and to compare gene expression between experimental groups. The Wald test was used to estimate p-values and log2 fold changes.

### Gene-set enrichment analysis

Gene ontology analysis was performed on statistically significant genes (p-adj < 0.05) with an absolute log fold change > 1. Briefly, the list of up- or down-regulated gene identifiers was uploaded separately onto Metascape and the results downloaded. Represented GO terms and graphics were adapted for clarity. For GSEA (Gene Set Enrichment Analysis), the normalized gene count matrix was used and zebrafish gene symbols were converted to their Human orthologs using the biomaRt package in R. Expression data file (.gct) and Phenotype data file (.cls) were generated according to GSEA guidelines. Files were uploaded into the GSEA software and analyzed with the following parameters:

Gene-set enrichment analysis (GSEA) was performed with the fgsea package. Genes were ranked in decreasing order by the DESeq2 Wald statistic. Zebrafish gene sets were obtained with msigdbr (*Danio rerio*) for the MSigDB Hallmark collection and the Gene Ontology Biological Process subcollection. Enrichment was computed with gene-set size limits of 15 to 500 genes and a fixed random seed; nominal P values were adjusted by the Benjamini-Hochberg method, and gene sets with an adjusted P value < 0.05 were considered significant. Normalized enrichment scores (NES) are reported. To reduce redundancy among Gene Ontology terms, near-duplicate terms with more than 50% gene overlap were collapsed, retaining the most significant representative. Running enrichment-score plots were generated from the fgsea enrichment data. As orthogonal collections, Reactome and KEGG gene sets were tested on the same Wald-statistic ranking using ReactomePA (gsePathway, organism “zebrafish”) and clusterProfiler (gseKEGG, organism “dre”), after mapping Ensembl identifiers to Entrez identifiers with org.Dr.eg.db (gene-set size 10 to 500).

### Immune and metabolic gene modules

The immune and metabolic gene sets used for the heatmaps were curated as the union of (i) the relevant Gene Ontology Biological Process term together with all of its descendant terms, mapped to zebrafish genes through org.Dr.eg.db, and (ii) manually curated zebrafish gene families, added to recover paralogues that are under-annotated in Gene Ontology. Metabolic genes were further assigned to modules (glycolysis and pyruvate metabolism, the tricarboxylic acid cycle, oxidative phosphorylation, fatty acid oxidation, mitochondrial translation, and creatine/phosphagen metabolism); genes belonging to more than one module were assigned to a single module by priority.

### Volcano plot and Heatmaps

Differential-expression results were displayed as a volcano plot (ggplot2), highlighting genes with an adjusted P value < 0.05 and an absolute log2 fold change ≥ 0.5; selected immune and metabolic genes were labeled with ggrepel. *lrrk1* was omitted from the plot for axis scaling only and was retained in all statistical analyses.

Heatmaps were generated with pheatmap from the blind VST matrix restricted to the selected genes. Expression values were converted to per-gene z-scores (row scaling) and displayed on a blue-white-red color scale saturating at ±2. Genes were ordered by module rather than by Euclidean clustering. Unnamed clones and a small number of ambiguous identifiers were excluded from the displayed metabolic heatmap.

### Quantification and Statistical Analysis

Sample sizes were chosen based on previous publications and are indicated in each figure or figure legend. No animal or sample was excluded from the analysis unless the animal died during the procedure. For proliferation and fibrosis experiments samples were assigned a numerical code to de-identify them as to experimental condition. These quantifications were done blinded. Other experiments were not randomized, and the investigators were not blinded to allocation during experiments and outcome assessment. All statistical values are displayed as mean ± standard deviation. Sample sizes, statistical test and *P* values are indicated in the figures or figure legends. Data distribution was determined before using parametric or non-parametric statistical test. Statistical significance was assigned at *P* < 0.05. All statistical tests were performed using Prism 9 software.

### Data availability

Raw and processed sequencing files are available on the Gene Expression Omnibus database under accession number GSE341742. Unique biological materials generated in this study (expression vectors and zebrafish lines) are available from the corresponding authors upon reasonable request.

## Notes

### Competing Interest Statement

The authors have declared no competing interest.

## REFERENCES

1. Poss KD, Wilson LG, Keating MT. Heart regeneration in zebrafish. Science. 2002;298:2188–2190. doi: 10.1126/science.1077857

2. Kikuchi K, Holdway JE, Werdich AA, Anderson RM, Fang Y, Egnaczyk GF, Evans T, Macrae CA, Stainier DY, Poss KD. Primary contribution to zebrafish heart regeneration by gata4(+) cardiomyocytes. Nature. 2010;464:601–605. doi: 10.1038/nature08804

3. Porrello ER, Mahmoud AI, Simpson E, Hill JA, Richardson JA, Olson EN, Sadek HA. Transient regenerative potential of the neonatal mouse heart. Science. 2011;331:1078–1080. doi: 10.1126/science.1200708

4. Gonzalez-Rosa JM, Sharpe M, Field D, Soonpaa MH, Field LJ, Burns CE, Burns CG. Myocardial Polyploidization Creates a Barrier to Heart Regeneration in Zebrafish. Dev Cell. 2018;44:433–446 e437. doi: 10.1016/j.devcel.2018.01.021

5. Frangogiannis NG. The inflammatory response in myocardial injury, repair, and remodelling. Nat Rev Cardiol. 2014;11:255–265. doi: 10.1038/nrcardio.2014.28

6. Steffens S, Van Linthout S, Sluijter JPG, Tocchetti CG, Thum T, Madonna R. Stimulating pro-reparative immune responses to prevent adverse cardiac remodelling: consensus document from the joint 2019 meeting of the ESC Working Groups of cellular biology of the heart and myocardial function. Cardiovasc Res. 2020;116:1850–1862. doi: 10.1093/cvr/cvaa137

7. Salimova E, Nowak KJ, Estrada AC, Furtado MB, McNamara E, Nguyen Q, Balmer L, Preuss C, Holmes JW, Ramialison M, et al. Variable outcomes of human heart attack recapitulated in genetically diverse mice. NPJ Regen Med. 2019;4:5. doi: 10.1038/s41536-019-0067-6

8. Patterson M, Barske L, Van Handel B, Rau CD, Gan P, Sharma A, Parikh S, Denholtz M, Huang Y, Yamaguchi Y, et al. Frequency of mononuclear diploid cardiomyocytes underlies natural variation in heart regeneration. Nat Genet. 2017;49:1346–1353. doi: 10.1038/ng.3929

9. Colak D, Kaya N, Al-Zahrani J, Al Bakheet A, Muiya P, Andres E, Quackenbush J, Dzimiri N. Left ventricular global transcriptional profiling in human end-stage dilated cardiomyopathy. Genomics. 2009;94:20–31. doi: 10.1016/j.ygeno.2009.03.003

10. Chaffin M, Papangeli I, Simonson B, Akkad AD, Hill MC, Arduini A, Fleming SJ, Melanson M, Hayat S, Kost-Alimova M, et al. Single-nucleus profiling of human dilated and hypertrophic cardiomyopathy. Nature. 2022. doi: 10.1038/s41586-022-04817-8

11. Vagnozzi RJ, Gatto GJ, Jr., Kallander LS, Hoffman NE, Mallilankaraman K, Ballard VL, Lawhorn BG, Stoy P, Philp J, Graves AP, et al. Inhibition of the cardiomyocyte-specific kinase TNNI3K limits oxidative stress, injury, and adverse remodeling in the ischemic heart. Sci Transl Med. 2013;5:207ra141. doi: 10.1126/scitranslmed.3006479

12. Theis JL, Zimmermann MT, Larsen BT, Rybakova IN, Long PA, Evans JM, Middha S, de Andrade M, Moss RL, Wieben ED, et al. TNNI3K mutation in familial syndrome of conduction system disease, atrial tachyarrhythmia and dilated cardiomyopathy. Hum Mol Genet. 2014;23:5793–5804. doi: 10.1093/hmg/ddu297

13. Xi Y, Honeywell C, Zhang D, Schwartzentruber J, Beaulieu CL, Tetreault M, Hartley T, Marton J, Vidal SM, Majewski J, et al. Whole exome sequencing identifies the TNNI3K gene as a cause of familial conduction system disease and congenital junctional ectopic tachycardia. Int J Cardiol. 2015;185:114–116. doi: 10.1016/j.ijcard.2015.03.130

14. Fan LL, Huang H, Jin JY, Li JJ, Chen YQ, Zhao SP, Xiang R. Whole exome sequencing identifies a novel mutation (c.333 + 2T > C) of TNNI3K in a Chinese family with dilated cardiomyopathy and cardiac conduction disease. Gene. 2018;648:63–67. doi: 10.1016/j.gene.2018.01.055

15. Ramzan S, Tennstedt S, Tariq M, Khan S, Noor Ul Ayan H, Ali A, Munz M, Thiele H, Korejo AA, Mughal AR, et al. A Novel Missense Mutation in TNNI3K Causes Recessively Inherited Cardiac Conduction Disease in a Consanguineous Pakistani Family. Genes (Basel). 2021;12. doi: 10.3390/genes12081282

16. Pham C, Andrzejczyk K, Jurgens SJ, Lekanne Deprez R, Palm KCA, Vermeer AMC, Nijman J, Christiaans I, Barge-Schaapveld D, van Dessel P, et al. Genetic Burden of TNNI3K in Diagnostic Testing of Patients With Dilated Cardiomyopathy and Supraventricular Arrhythmias. Circ Genom Precis Med. 2023;16:328–336. doi: 10.1161/CIRCGEN.122.003975

17. Tang H, Xiao K, Mao L, Rockman HA, Marchuk DA. Overexpression of TNNI3K, a cardiac-specific MAPKKK, promotes cardiac dysfunction. J Mol Cell Cardiol. 2013;54:101–111. doi: 10.1016/j.yjmcc.2012.10.004

18. Wheeler FC, Tang H, Marks OA, Hadnott TN, Chu PL, Mao L, Rockman HA, Marchuk DA. Tnni3k modifies disease progression in murine models of cardiomyopathy. PLoS Genet. 2009;5:e1000647. doi: 10.1371/journal.pgen.1000647

19. Reuter SP, Soonpaa MH, Field D, Simpson E, Rubart-von der Lohe M, Lee HK, Sridhar A, Ware SM, Green N, Li X, et al. Cardiac Troponin I-Interacting Kinase Affects Cardiomyocyte S-Phase Activity but Not Cardiomyocyte Proliferation. Circulation. 2023;147:142–153. doi: 10.1161/CIRCULATIONAHA.122.061130

20. Purdy AL, Swift SK, Sucov HM, Patterson M. Tnni3k influences cardiomyocyte S-phase activity and proliferation. J Mol Cell Cardiol. 2023;183:22–26. doi: 10.1016/j.yjmcc.2023.08.004

21. Gan P, Baicu C, Watanabe H, Wang K, Tao G, Judge DP, Zile MR, Makita T, Mukherjee R, Sucov HM. The prevalent I686T human variant and loss-of-function mutations in the cardiomyocyte-specific kinase gene TNNI3K cause adverse contractility and concentric remodeling in mice. Hum Mol Genet. 2021;29:3504– 3515. doi: 10.1093/hmg/ddaa234

22. Gonzalez-Rosa JM, Martin V, Peralta M, Torres M, Mercader N. Extensive scar formation and regression during heart regeneration after cryoinjury in zebrafish. Development. 2011;138:1663–1674. doi: 10.1242/dev.060897

23. Jopling C, Sleep E, Raya M, Marti M, Raya A, Izpisua Belmonte JC. Zebrafish heart regeneration occurs by cardiomyocyte dedifferentiation and proliferation. Nature. 2010;464:606–609. doi: 10.1038/nature08899

24. Pfefferli C, Jazwinska A. The careg element reveals a common regulation of regeneration in the zebrafish myocardium and fin. Nat Commun. 2017;8:15151. doi: 10.1038/ncomms15151

25. Feng Y, Cao HQ, Liu Z, Ding JF, Meng XM. Identification of the dual specificity and the functional domains of the cardiac-specific protein kinase TNNI3K. Gen Physiol Biophys. 2007;26:104–109.

26. Wang X, Wang J, Su M, Wang C, Chen J, Wang H, Song L, Zou Y, Zhang L, Zhang Y, et al. TNNI3K, a cardiac-specific kinase, promotes physiological cardiac hypertrophy in transgenic mice. PLoS One. 2013;8:e58570. doi: 10.1371/journal.pone.0058570

27. Li H, Trager LE, Liu X, Hastings MH, Xiao C, Guerra J, To S, Li G, Yeri A, Rodosthenous R, et al. lncExACT1 and DCHS2 Regulate Physiological and Pathological Cardiac Growth. Circulation. 2022;145:1218–1233. doi: 10.1161/CIRCULATIONAHA.121.056850

28. Lekkos K, Hu Z, Nguyen PD, Honkoop H, Sengul E, Alonaizan R, Koth J, Ying J, Lemieux ME, Kenward A, et al. Oxidative phosphorylation is required for cardiomyocyte re-differentiation and long-term fish heart regeneration. Nat Cardiovasc Res. 2025;4:1363–1380. doi: 10.1038/s44161-025-00718-x

29. Willeford A, Suetomi T, Nickle A, Hoffman HM, Miyamoto S, Heller Brown J. CaMKIIdelta-mediated inflammatory gene expression and inflammasome activation in cardiomyocytes initiate inflammation and induce fibrosis. JCI Insight. 2018;3. doi: 10.1172/jci.insight.97054

30. Simoes FC, Cahill TJ, Kenyon A, Gavriouchkina D, Vieira JM, Sun X, Pezzolla D, Ravaud C, Masmanian E, Weinberger M, et al. Macrophages directly contribute collagen to scar formation during zebrafish heart regeneration and mouse heart repair. Nat Commun. 2020;11:600. doi: 10.1038/s41467-019-14263-2

31. Bevan L, Lim ZW, Venkatesh B, Riley PR, Martin P, Richardson RJ. Specific macrophage populations promote both cardiac scar deposition and subsequent resolution in adult zebrafish. Cardiovasc Res. 2020;116:1357–1371. doi: 10.1093/cvr/cvz221

32. de Preux Charles AS, Bise T, Baier F, Marro J, Jazwinska A. Distinct effects of inflammation on preconditioning and regeneration of the adult zebrafish heart. Open Biol. 2016;6. doi: 10.1098/rsob.160102

33. Lai SL, Marin-Juez R, Moura PL, Kuenne C, Lai JKH, Tsedeke AT, Guenther S, Looso M, Stainier DY. Reciprocal analyses in zebrafish and medaka reveal that harnessing the immune response promotes cardiac regeneration. Elife. 2017;6. doi: 10.7554/eLife.25605

34. Rihan M, Zangi L, Magadum A. Pyruvate Kinase M2 Role in Cardiovascular Repair. Cells. 2025;14. doi: 10.3390/cells14201623

35. Stone OA, El-Brolosy M, Wilhelm K, Liu X, Romao AM, Grillo E, Lai JKH, Gunther S, Jeratsch S, Kuenne C, et al. Loss of pyruvate kinase M2 limits growth and triggers innate immune signaling in endothelial cells. Nat Commun. 2018;9:4077. doi: 10.1038/s41467-018-06406-8

36. Keeley S, Fernandez-Lajarin M, Bergemann D, John N, Parrott L, Andrea BE, Gonzalez-Rosa JM. Rapid and robust generation of cardiomyocyte-specific crispants in zebrafish using the cardiodeleter system. Cell Rep Methods. 2025;5:101003. doi: 10.1016/j.crmeth.2025.101003

37. Lai ZF, Chen YZ, Feng LP, Meng XM, Ding JF, Wang LY, Ye J, Li P, Cheng XS, Kitamoto Y, et al. Overexpression of TNNI3K, a cardiac-specific MAP kinase, promotes P19CL6-derived cardiac myogenesis and prevents myocardial infarction-induced injury. Am J Physiol Heart Circ Physiol. 2008;295:H708–716. doi: 10.1152/ajpheart.00252.2008

38. Burke MA, Chang S, Wakimoto H, Gorham JM, Conner DA, Christodoulou DC, Parfenov MG, DePalma SR, Eminaga S, Konno T, et al. Molecular profiling of dilated cardiomyopathy that progresses to heart failure. JCI Insight. 2016;1. doi: 10.1172/jci.insight.86898

39. Teekakirikul P, Eminaga S, Toka O, Alcalai R, Wang L, Wakimoto H, Nayor M, Konno T, Gorham JM, Wolf CM, et al. Cardiac fibrosis in mice with hypertrophic cardiomyopathy is mediated by non-myocyte proliferation and requires Tgf-beta. J Clin Invest. 2010;120:3520–3529. doi: 10.1172/JCI42028

40. Pan YA, Freundlich T, Weissman TA, Schoppik D, Wang XC, Zimmerman S, Ciruna B, Sanes JR, Lichtman JW, Schier AF. Zebrabow: multispectral cell labeling for cell tracing and lineage analysis in zebrafish. Development. 2013;140:2835–2846. doi: 10.1242/dev.094631

41. Moreno-Mateos MA, Vejnar CE, Beaudoin JD, Fernandez JP, Mis EK, Khokha MK, Giraldez AJ. CRISPRscan: designing highly efficient sgRNAs for CRISPR-Cas9 targeting in vivo. Nat Methods. 2015;12:982–988. doi: 10.1038/nmeth.3543

42. Shah AN, Davey CF, Whitebirch AC, Miller AC, Moens CB. Rapid reverse genetic screening using CRISPR in zebrafish. Nat Methods. 2015;12:535–540. doi: 10.1038/nmeth.3360

43. Vejnar CE, Moreno-Mateos MA, Cifuentes D, Bazzini AA, Giraldez AJ. Optimized CRISPR-Cas9 System for Genome Editing in Zebrafish. Cold Spring Harb Protoc. 2016;2016. doi: 10.1101/pdb.prot086850

44. Kanther M, Sun X, Muhlbauer M, Mackey LC, Flynn EJ, 3rd, Bagnat M, Jobin C, Rawls JF. Microbial colonization induces dynamic temporal and spatial patterns of NF-kappaB activation in the zebrafish digestive tract. Gastroenterology. 2011;141:197–207. doi: 10.1053/j.gastro.2011.03.042

45. Wei KH, Lin IT, Chowdhury K, Lim KL, Liu KT, Ko TM, Chang YM, Yang KC, Lai SB. Comparative single-cell profiling reveals distinct cardiac resident macrophages essential for zebrafish heart regeneration. Elife. 2023;12. doi: 10.7554/eLife.84679

46. Gonzalez-Rosa JM, Mercader N. Cryoinjury as a myocardial infarction model for the study of cardiac regeneration in the zebrafish. Nat Protoc. 2012;7:782–788. doi: 10.1038/nprot.2012.025.

