## Supplemental Figures for "High levels of the cardiomyocyte-specific kinase Tnni3k impair zebrafish heart regeneration by driving chronic myocardial inflammation"

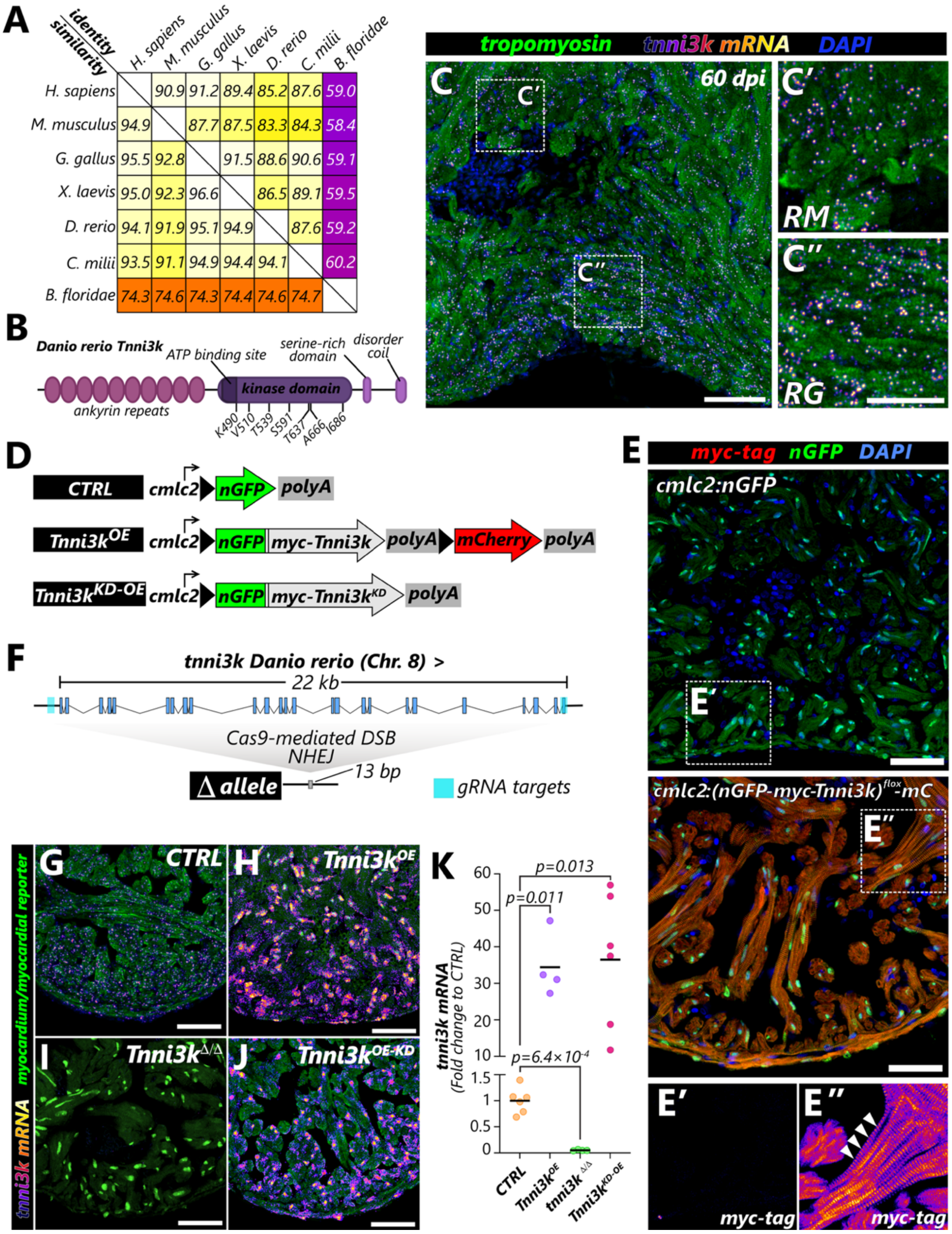


**Supplementary Figure 1**. **Tnni3k is evolutionarily conserved, upregulated after injury, highly expressed in the Tnni3k$^{OE}$ zebrafish line, and absent in the *Tnni3k* mutant line.**

**(A)** Conservation degree of Tnni3k protein sequence among species represented as the percentage of similarity and identity.

**(B)** Schematic representation of *Danio rerio* Tnni3k protein main domains. Key residues previously described as critical for the kinase function are indicated.

**(C)** *tnni3k* RNAscope images of representative ventricular sections 60 days after cryoinjury (dpi). Magnification of boxed areas is shown in C' and C''.

**(D)** DNA constructs used to drive cardiomyocyte-specific expression of GFP (*CTRL*), *tnni3k* (*Tnni3k*$^{OE}$), or a kinase-dead *tnni3k* (*Tnni3k*$^{KD\text{-}OE}$).

**(E)** Immunofluorescence images from control and *Tnni3k*$^{OE}$ hearts detecting nuclear GFP and myc-tagged Tnni3k. Magnified boxed areas show myc-tag signal of control (E') and *Tnni3k*$^{OE}$ hearts (E").

**(F)** CRISPR/Cas9 strategy used to delete the 22-kb *tnni3k* locus on chromosome 8 (gRNA target sites in cyan).

**(G-J)** *In situ* hybridization (RNAscope) images of heart sections of *CTRL* (G)*,* Tnni3k$^{OE}$(H), *tnni3k*$^{\Delta/\Delta}$ (I) and Tnni3k$^{KD\text{-}OE}$ (J) animal whole hearts to visualize *Tnni3k* mRNA.

**(K)** *tnni3k* mRNA quantification by RT-qPCR in *CTRL,* Tnni3k$^{OE}$, *tnni3k*$^{\Delta/\Delta}$ and Tnni3k$^{KD\text{-}OE}$ animal whole hearts. *rps11* RNA was used to normalize *tnni3k* mRNA content in each sample. *p* values: Brown-Forsythe and Welch ANOVA followed by Dunnett's multiple comparison test. 3 pooled hearts/sample.

RG, regenerated myocardium; RM, remote myocardium

Scale bars, 50 μm.

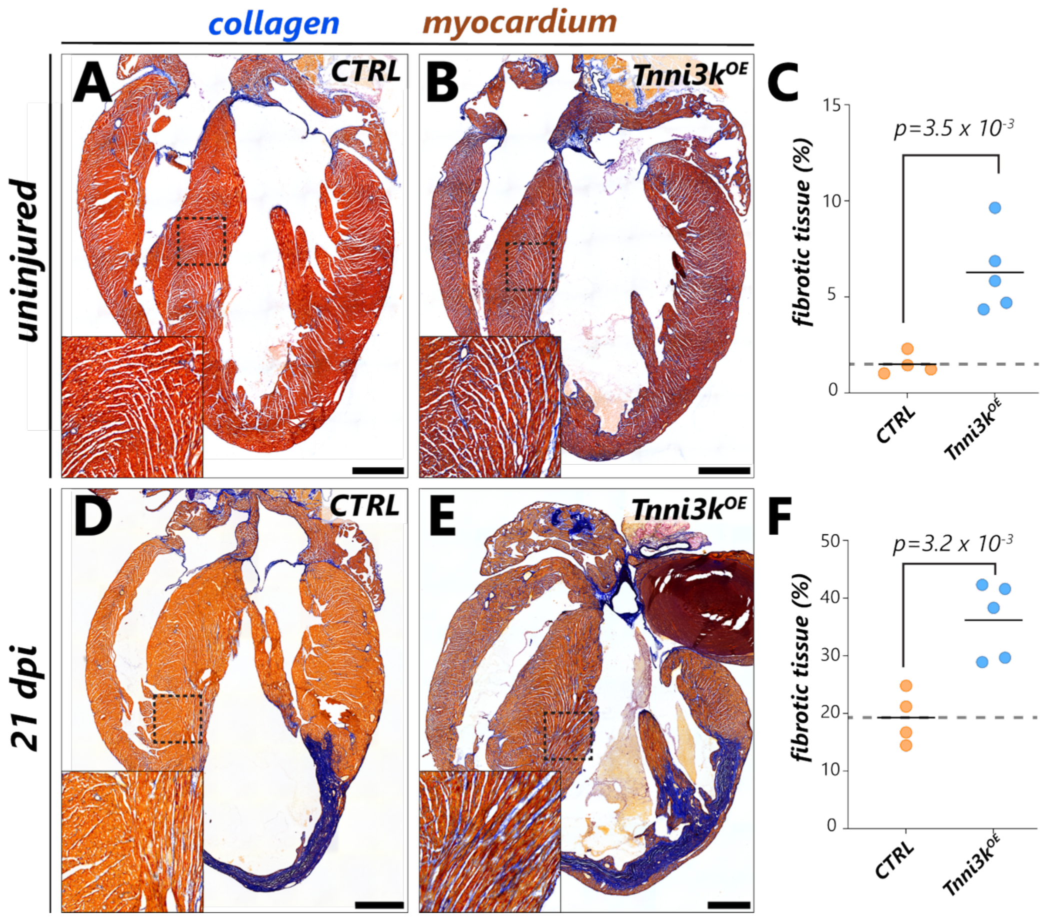


**<u>Supplementary Figure 2</u>. Tnni3k overexpression increases fibrosis in the homeostatic mouse heart and after injury in a model of myocardial infarction.**

**(A, B)** Representative images of AFOG-stained cryosections of uninjured hearts from wild-type DAB (A) and Tnni3k overexpressing (B) mouse lines. Magnifications of the boxed areas are shown in the bottom-left corner.

**(C)** Fibrosis quantification relative to the entire area of the chambers for the indicated cohorts in uninjured hearts. 3 sections analyzed per sample, n (CTRL) =4, n(Tnni3k$^{OE}$) = 5. Black line, average. *p*-values: two-tailed unpaired *t-test.*

**(D, E)** AFOG-stained cryosections of hearts from wild-type DAB (D) and Tnni3k overexpressing (E) mouse lines 21 days after coronary artery ligation (21 dpi). Magnification of the boxed areas is also shown**.**

**(F)** Fibrosis quantification relative to the entire area of the chambers for the indicated cohorts in hearts at 21 dpi. 3 sections analyzed per sample, n (CTRL) =4, n(Tnni3k$^{OE}$) = 5. Black line, average. *p* values: two-tailed unpaired *t*-test*.* Scale bar, 1 mm.

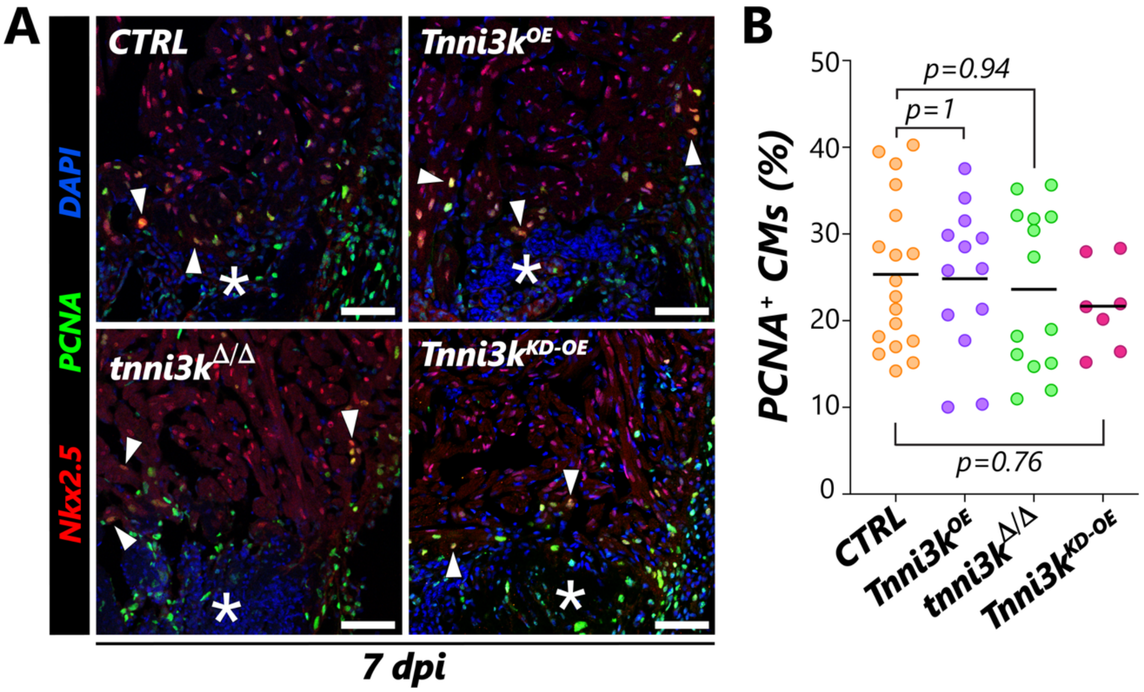


**<u>Supplementary Figure 3</u>. Tnni3k overexpression does not affect cardiomyocyte cell cycle entry at 7 days post-cryoinjury.**

**(A)** Heart sections from *CTRL*, *Tnni3k*$^{OE}$, *tnni3k*$^{\Delta/\Delta}$ and *Tnni3k*$^{KD\text{-}OE}$ fish at 7 days post-cryoinjury (dpi) immunostained for Nkx2.5 (cardiomyocyte nuclei) and PCNA (proliferating cell nuclear antigen, cycling cells). Arrowheads, Nkx2.5$^+$ PCNA$^+$ cardiomyocytes; asterisks, injury zone.

**(B)** Quantification of cycling cardiomyocytes as the percentage of cardiomyocytes positive for the PCNA marker (Nkx2.5$^+$, PCNA$^+$) relative to the total number of cardiomyocytes (Nkx2.5$^+$) within a defined area (150 μm x 400 μm) in the injury border zone. *p*-values: one-way ANOVA followed by Tukey's multiple comparison test. 3 sections analyzed/sample.

Scale bars, 50 μm.

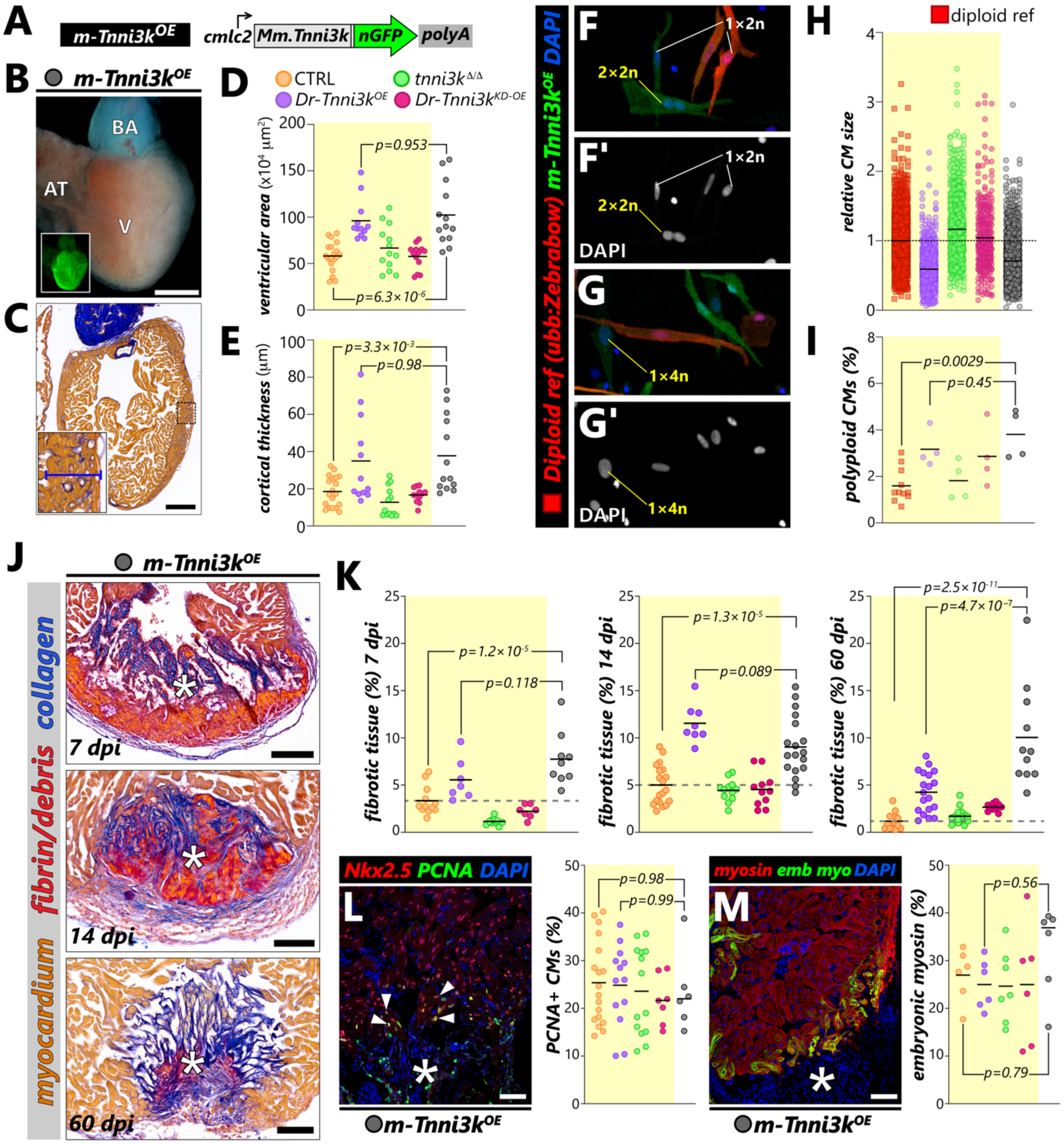


**Supplementary Figure 4. Zebrafish overexpressing mouse *Tnni3k* reproduce the ventricular enlargement, modest polyploidization, and impaired heart regeneration of the zebrafish *tnni3k* overexpressing line.**

**(A)** DNA construct used to generate the *mouse-Tnni3k*$^{OE}$ line, driving cardiomyocyte-specific expression of *Tnni3k* from *Mus musculus* and nuclear GFP.

**(B)** Representative image from *mouse-Tnni3k*$^{OE}$ zebrafish whole heart. GFP fluorescence from the transgene is shown in the bottom-left corner.

**(C)** Histologic section of a *mouse-Tnni3k*$^{OE}$ heart stained with Acid Fuchsin Orange G (AFOG) technique. Magnification of the boxed area is shown in the bottom-left corner.

**(D, E)** Quantification of ventricular area (D) and cortical thickness (E) on AFOG-stained tissue sections from the indicated cohorts. Data already presented in **Figure 2** are shaded in yellow for reference.

**(F-G')** Representative images from cardiomyocyte dissociation of diploid reference (*ubb:Zebrabow)* and mouse-*Tnni3k*$^{OE}$ ventricles stained with the DNA dye DAPI. DAPI channel is shown in (F') and (G') to illustrate different ploidy classes.

**(H, I)** Quantification of cardiomyocyte size (H) and frequency of polyploid cardiomyocytes (I) from ventricular dissociations of the indicated cohorts (3 pooled ventricles per sample, ~1,000 CMs analyzed per sample). Yellow shading indicates data previously shown in **Figure 2**. *p* values: One-way ANOVA, followed by Tukey's multiple comparison test.

**(J)** Acid Fuchsin Orange G (AFOG) stained heart sections from *mouse-Tnni3k*$^{OE}$ animals at 7-, 14- and 60 days post-cryoinjury (dpi). Asterisk, injury zone.

**(K)** Quantification of fibrotic area relative to ventricular area in hearts from the indicated cohorts at 7-, 14- and 60dpi. Yellow shading indicates data previously shown in **Figure 2**. *p* values: One-way ANOVA, followed by Tukey's multiple comparison test.

**(L)** Heart section from *mouse-Tnni3k*$^{OE}$ fish at 7dpi immunostained for Nkx2.5 (cardiomyocyte nuclei) and PCNA (proliferating cell nuclear antigen, cycling cells), and quantification of cycling cardiomyocytes (Nkx2.5$^+$, PCNA$^+$) relative to the total number of cardiomyocytes (Nkx2.5$^+$) within a defined area in the injury border zone. Arrowheads, Nkx2.5$^+$ PCNA$^+$ cardiomyocytes; asterisks, injury zone. The data previously shown in **Figure 2 are** highlighted in yellow. *p*-values: one-way ANOVA followed by Tukey's multiple comparison test. 3 sections analyzed/sample.

**(M)** Heart section images at 7dpi immunostained to detect cardiomyocyte myosin heavy chain and the embryonic form of myosin heavy chain, and quantification of embryonic myosin heavy chain expressing myocardium relative to myocardium area. Asterisks, injury zone. Yellow shading indicates data previously shown in **Figure 2**. *p* values: one-way ANOVA followed by Tukey's multiple comparison test. 3 sections analyzed/sample.

Scale bars: 1 mm (B-C), 200 μm (J), 100 μm (L, M).

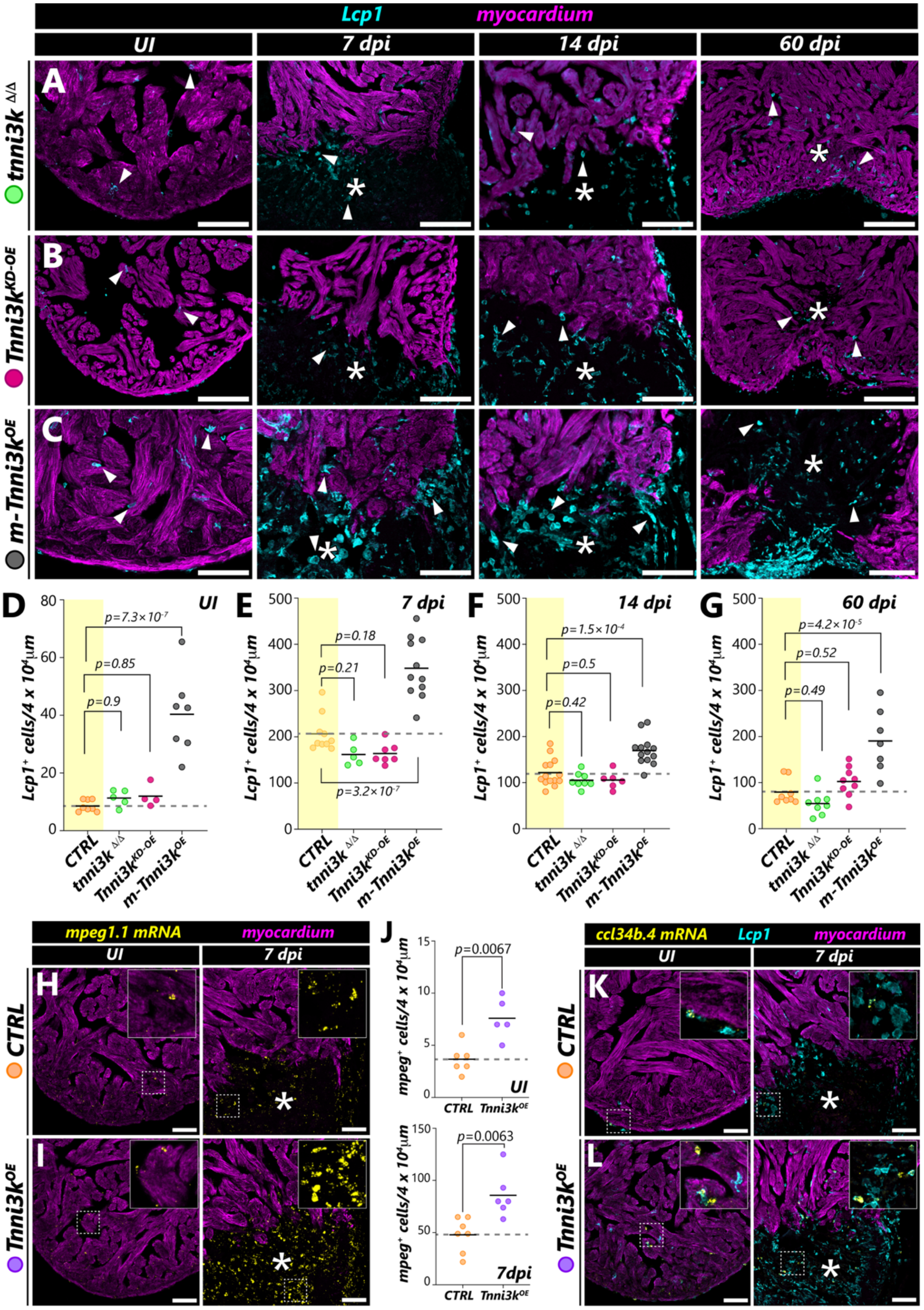


**Supplementary Figure 5.** **Only wild type Tnni3k overexpression triggers proinflammatory states.**

**(A-C)** Representative heart sections from *tnni3k*$^{\Delta/\Delta}$ (A), *Tnni3k*$^{KD\text{-}OE}$ (B) and m-*Tnni3k*$^{OE}$ (C) of uninjured (UI) hearts and at 7, 14, and 60 days post-cryoinjury (dpi) stained for leukocyte (Lcp1$^{+}$) and myocardium (myosin heavy chain) detection. Arrowheads, leukocytes. Asterisks, injured area.

**(D-G)** Quantification of leukocyte number in hearts from the indicated cohorts in uninjured (UI) conditions (D) and at 7 (E), 14 (F) and 60 (G) dpi. n (*UI*) = 8, 5, 4, 7; n (7 dpi) =10, 5, 7, 11; n (14 dpi)= 14, 8, 6, 13; n (60 dpi) = 8, 8, 9, 7. The data previously shown in **Figure 6** are highlighted in yellow. *p* values: one-way ANOVA, followed by Tukey's multiple comparison test. 3 sections analyzed/sample.

**(H-I)** RNAscope *in situ* hybridization and immunofluorescence images for macrophage marker *mpeg1.1* RNA and myosin heavy chain protein co-detection in *CTRL* (H) and *Tnni3k*$^{OE}$ (I) uninjured hearts and at 7 dpi. Magnifications of the boxed areas are shown in the upper-right corner. Asterisks, injured area.

**(J)** Quantification of *mpeg1.1*$^{+}$ cells (macrophages) in defined areas containing the injury zone, for the indicated conditions and cohorts. *p*-value*s*: two-tailed unpaired *t-test*

**(K, L)** Representative images of *ccl34b.4* RNAscope *in situ* hybridization and myosin heavy chain immunofluorescence in *CTRL* (K) and *Tnni3k*$^{OE}$ (L) uninjured hearts and at 7 dpi. Magnifications of the boxed areas are shown in the upper-right corner. Asterisks, injured area.

Scale bars, 50 μm.

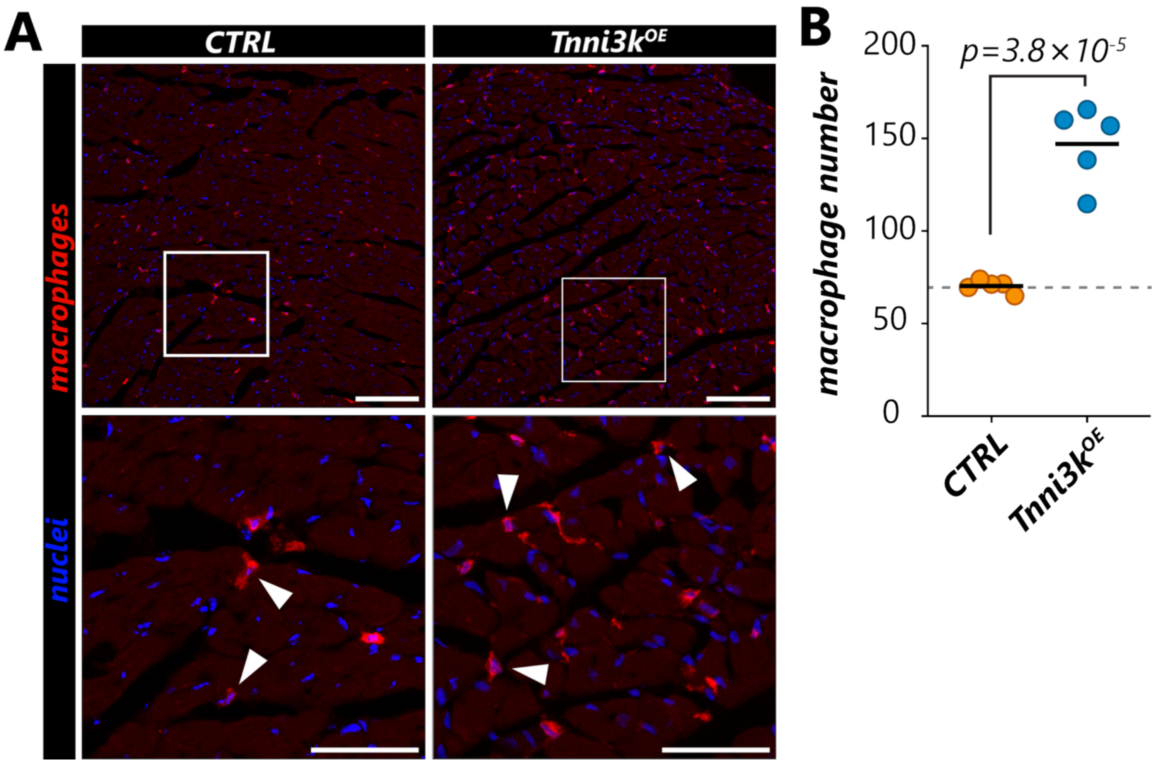


**<u>Supplementary Figure 6.</u>** ***TNNI3K*** **overexpression increases macrophage number in the homeostatic mouse heart.**

**(A)** Representative immunofluorescence images of mouse heart cryosections from wild-type DAB (*CTRL*) and Tnni3k overexpressing (*Tnni3k*$^{OE}$) mouse lines detecting macrophages (F4/80 pan-macrophage marker positive cells). Magnification of the boxed areas is shown at the bottom. Arrowheads, macrophages.

**(B)** Quantification of macrophage number in hearts from the indicated cohorts. Dashed line, average. *p*-value: two-tailed unpaired *t-test*. 3 sections analyzed/sample; 3 areas analyzed/section.

Scale bars, 100μm (top); 50μm (bottom).
